# The extracellular matrix supports cancer cell growth under amino acid starvation by promoting tyrosine catabolism

**DOI:** 10.1101/2021.06.09.447520

**Authors:** Mona Nazemi, Bian Yanes, Montserrat Llanses Martinez, Heather Walker, Frederic Bard, Elena Rainero

## Abstract

Breast and pancreatic tumours are embedded in a collagen I-rich extracellular matrix (ECM) network, where nutrients are scarce due to limited blood flow and elevated tumour growth. Metabolic adaptation is required for cancer cells to endure these conditions. Here, we demonstrated that the presence of ECM supported the growth of invasive breast cancer cells, but not non-transformed mammary epithelial cells, and pancreatic cancer cells under amino acid starvation, through a mechanism that required macropinocytosis-dependent ECM uptake. Importantly, we showed that this behaviour was acquired during carcinoma progression. ECM internalisation, followed by lysosomal degradation, contributed to the upregulation of the intracellular levels of several amino acids, most notably tyrosine and phenylalanine. This resulted in elevated tyrosine catabolism on ECM under starvation, leading to increased fumarate levels, potentially feeding into the tricarboxylic acid cycle. Interestingly, this pathway was required for ECM-dependent cell growth under amino acid starvation, as the knockdown of p-hydroxyphenylpyruvate hydroxylase-like protein (HPDL), the third enzyme of the pathway, opposed cell growth on ECM in both 2D and 3D systems, without affecting cell proliferation on plastic. Finally, high HPDL expression correlated with poor prognosis in breast and pancreatic cancer patients. Collectively, our results highlight that the ECM in the tumour microenvironment represents an alternative source of nutrients to support cancer cell growth, by regulating phenylalanine and tyrosine metabolism.

## Introduction

Breast cancer is the most common type of cancer among women. Breast cancer progression starts from benign hyperplasia of epithelial cells of the mammary duct, leading to atypical ductal hyperplasia and ductal carcinoma in situ (DCIS), where cancer cells are surrounded by an intact basement membrane (BM). Eventually, the cells acquire the ability to break through the BM, resulting in the invasive and metastatic cancer phenotype^1^. Pancreatic cancer has one of the poorest prognosis rates, with a 5-years survival rate of about 10%. Around 95% of pancreatic cancers are derived from exocrine cells and the most common form is pancreatic ductal adenocarcinoma (PDAC)^2^. A key feature of PDAC, both in primary and metastatic sites, is a highly desmoplastic stroma. This dense desmoplasia is composed of activated cancer-associated fibroblasts (CAFs) and collagen network, as well as immune cells^3^. In breast cancer, high breast tissue density is associated with a shift to malignancy and invasion^4, 5^. There is growing evidence indicating that the tumour microenvironment (TME) facilitates tumour growth and survival^6^. The TME consists of a variety of cell types, including stromal cells, and extracellular matrix (ECM). The ECM is a highly dynamic three-dimensional network of macromolecules, providing structural and mechanical support to the tissues while interacting with cells through different receptors^7, 8^.

Due to limited blood supply and elevated tumour growth rate, the TME is often deprived of nutrients, including glucose and amino acids^9^. One of the hallmarks of cancer is the reprogramming of energy metabolism, which provides metabolic flexibility allowing tumour cells to adapt to different nutrient conditions^10^. This includes the ability of cancer cells to benefit from alternative nutrient sources. Indeed, different studies showed that cancer cells benefit from extracellular proteins during food scarcity^11–13^. In Ras-driven PDAC cells, amino acid and glutamine starvation induced albumin and collagen internalisation respectively, followed by lysosomal degradation and amino acid extraction^9, 12, 14, 15^. Serum and growth factor starvation also stimulated normal mammary epithelial cells to internalise laminin, which resulted in an increase in cellular amino acid content^11^. These studies prompted us to investigate how the internalisation of different components of the ECM impinged on cancer cells’ metabolism and growth under amino acid starvation.

To determine whether the ECM surrounding breast cancer cells could provide a metabolically favourable microenvironment, here we assessed the effect of the presence of ECM on breast cancer cell growth and metabolism under glutamine and full amino acid starvation. We demonstrated that the growth of starved invasive breast cancer cells, but not normal mammary epithelial cells, was supported by the presence of different types of ECM. Indeed, Matrigel, collagen I and CAFs-generated cell derived matrix (CDM) promoted MDA-MB-231 cell division and enhanced mammalian target of rapamycin complex 1 (mTORC1) activity under amino acid starvation. Interestingly, this was dependent on ECM macropinocytosis and not focal adhesion signalling. Moreover, metabolomics analyses showed higher amino acid content and upregulation of phenylalanine and tyrosine metabolism during amino acid deprivation in cancer cells grown on ECM compared to plastic, leading to an increase in fumarate content. Importantly, the down-regulation of an enzyme in this pathway (HPDL) strongly reduced cell growth on ECM under amino acid starvation, without affecting cell growth on plastic. Interestingly, we demonstrated that the same pathway was also supporting the growth of PDAC cells under amino acid starvation. Moreover, high HPDL expression correlated with poor prognosis in breast and pancreatic cancer patients. Altogether, our data showed that ECM scavenging fuels cancer cell growth by promoting phenylalanine and tyrosine catabolism.

## Results

### The ECM supported invasive breast cancer cell growth under starvation

To assess whether the presence of ECM could promote the survival or growth of invasive breast cancer cells under nutrient deprivation, MDA-MB-231 cells were seeded under complete media, glutamine or amino acid deficiency and cell growth was quantified on collagen I, Matrigel and plastic. MDA-MB-231 cells showed significantly higher cell number and growth rate in the presence of collagen I and Matrigel compared to cells on plastic after eight days of starvation (**Figure 1A-C; Figure S1A-C**). Our data also indicated that, in the absence of starvation, cell growth was independent of the ECM, as there was no significant difference in cell number on ECM compared to plastic in complete media (**Figure S1D-F**). To test the effect of glutamine and amino acid deficiency on cell growth in a more physiological environment, MDA-MB-231 were seeded on cell-derived matrices (CDMs) generated by either normal breast fibroblasts (NFs) or cancer-associated fibroblasts (CAFs) extracted from breast tumours. CDMs are complex 3D matrices that recapitulate several features of native collagen-rich matrices^16^. Different studies highlighted that the CDM produced by CAFs has different structure and composition compared to the CDM derived from normal fibroblasts^17, 18^. Here, we showed that breast cancer cell growth was rescued by both NF and CAF generated CDM under glutamine starvation, but only CAF-CDM resulted in a significant increase in cell growth under amino acid deprivation (**Figure 1D,E**). These data indicate that CAF-CDM provided a more favourable environment for cancer cells’ growth compared to NF-CDM under amino acid deprivation. In parallel, to more closely reproduce the in vivo metabolic environment, we used a physiological medium, Plasmax. Plasmax is designed to contain metabolites and nutrients at the same concentration of human blood to avoid adverse effects of commercial media on cancer cells^19^. To simulate the TME starvation conditions, Plasmax was diluted with PBS. Cell growth data demonstrated that, while cells on plastic were not able to grow in 25 % Plasmax, the presence of collagen I, Matrigel and CAF-generated CDM resulted in a significant increase in cell numbers (**Figure 1F-H**). Moreover, the ECM had a small but significant effect in promoting cell growth in full Plasmax media as well (**Figure S1G**), suggesting that under more physiological conditions the ECM could support cell growth even in the absence of nutrient starvation. Taken together, these data indicated that the presence of ECM positively affected invasive breast cancer cell growth under nutrient depleted conditions.

**Figure 1.**
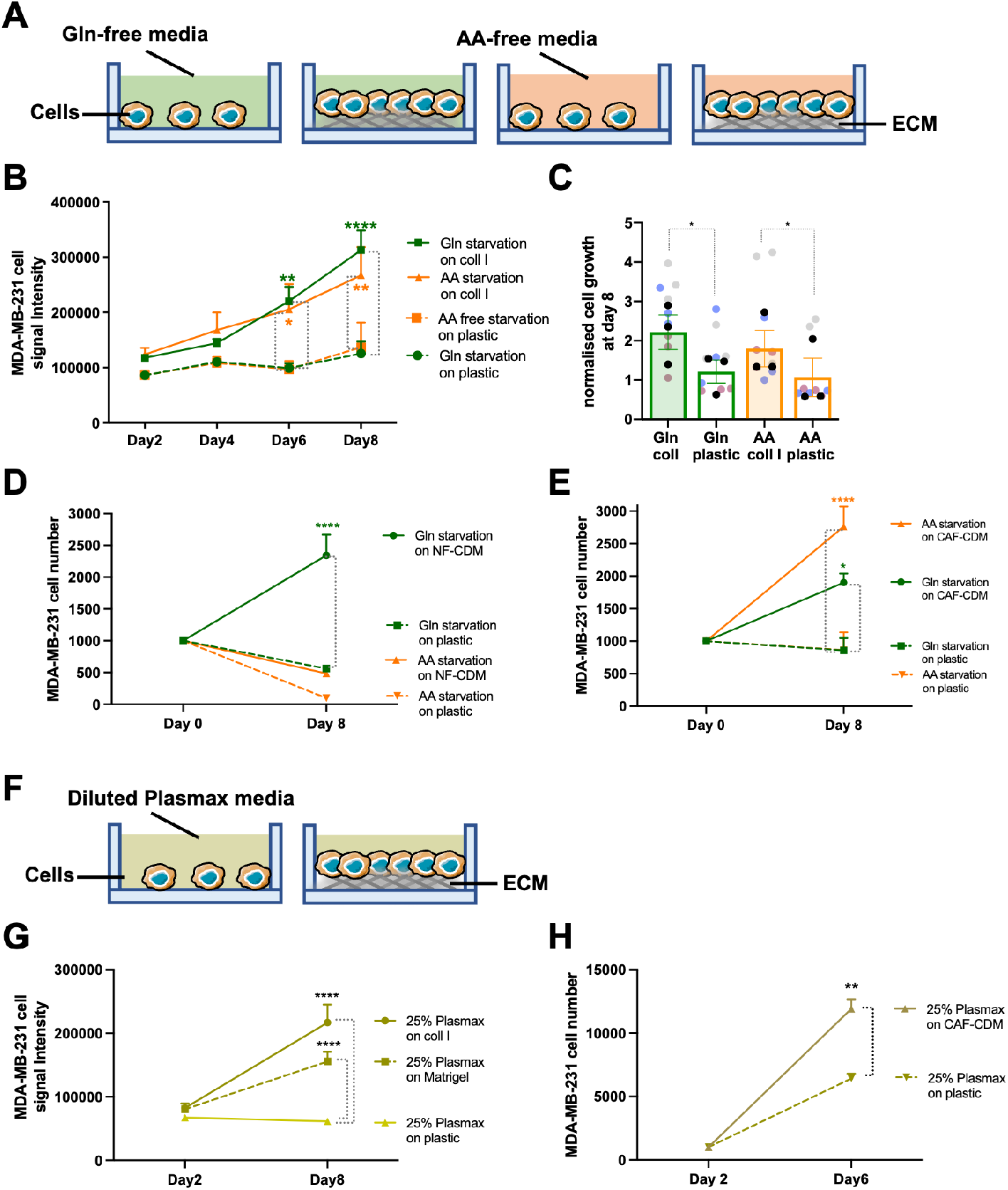
The ECM supported cell growth under starvation. (**A,F**) Schematic, cell proliferation assays. (**B,C**) MDA-MB-231 cells were seeded on plastic or 2mg/ml collagen I (coll I) for 8 days under glutamine (Gln) and amino acid (AA) starvation, fixed, stained with DRAQ5 and imaged with a Licor Odyssey system. Signal intensity was calculated by Image Studio Lite software. MDA-MB-231 cells were seeded on plastic, (**D**) NF-CDM or (**E**) CAF-CDM under Gln or AA starvation for 8 days, fixed and stained with Hoechst 33342. Images were collected by ImageXpress micro and analysed by MetaXpress software. MDA-MB-231 cells were seeded on plastic, 2mg/ml collagen I (coll I) or 3mg/ml Matrigel (**G**) or on CAF-CDM (**H**) in Plasmax media diluted 1:4 in PBS (25%) for 8 days, fixed and stained with DRAQ5 (**G**) or Hoechst 33342 (**H**) and quantified as above. Values are mean ± SEM from 3 independent experiments (the black dots in the bar graphs represent the mean of individual experiments). *p<0.05, **p<0.01, **** p<0.0001 (**B,D,E,G,H**) 2way ANOVA, Tukey’s multiple comparisons test. (**C**) Kruskal-Wallis, Dunn’s multiple comparisons test.

To assess whether ECM-dependent cell growth under starvation was a feature that was acquired during breast cancer progression, we took advantage of the MCF10 series of cells lines^20^. This includes normal mammary epithelial cells (MCF10A, **Figure 2A**), non-invasive ductal carcinoma in-situ breast cancer cells (MCF10A-DCIS, **Figure 2D**) and metastatic breast cancer cells (MCF10CA1, **Figure 2G**). The cells were seeded either on plastic, collagen I or Matrigel in glutamine or amino acid depleted media for eight days. In contrast to invasive MDA-MB-231 cells (Figure 1), MCF10A cell growth pattern on collagen I and Matrigel under starvation was similar to the one on plastic. Although cell number was higher on collagen I and Matrigel under amino acid starvation compared to plastic (**Figure 2B; Figure S2B**), there was no significant difference in the growth rate of MCF10A cells on either matrix (**Figure 2C; Figure S2C**). Similarly, our data showed no significant changes in MCF10A-DCIS cell growth on collagen I or Matrigel in amino acid depleted media compared to the plastic (**Figure 2E,F; Figure S2E,F**). Interestingly, while collagen I was not able to increase MCF10A-DCIS cell growth under glutamine deficiency (**Figure 2E,F**), cell number and growth rate at day eight post glutamine starvation were significantly higher on Matrigel compared to plastic (**Figure S2E,F**). Consistent with our observation in MDA-MB-231 cells, the growth of metastatic MCF10CA1 cells was significantly promoted by collagen I, CAF-CDM and Matrigel under glutamine or amino acid starvation (**Figure 2H-J; Figure S2H,I**). As before, the presence of ECM did not affect MCF10A, MCF10A-DCIS and MCF10CA1 cells under complete media, with the exception of Matrigel resulting in a small but significant increase in MCF10A cell growth (**Figure S2J-O**). These data suggest that the ability of using ECM to compensate for soluble nutrient starvation could have been gradually acquired during cancer progression and could be a defining feature of invasive growth in advanced metastatic breast cancer.

**Figure 2.**
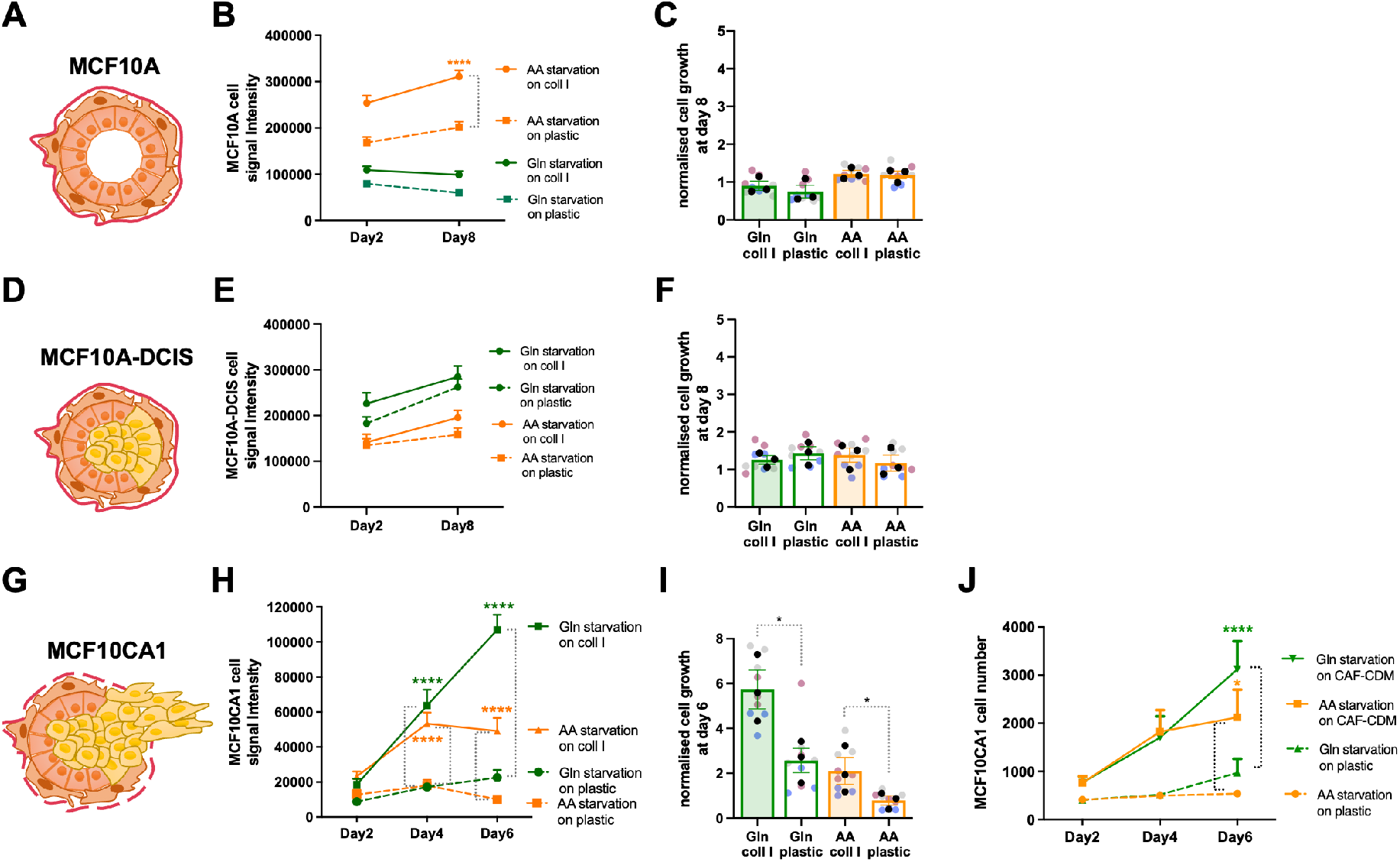
The ability to use the ECM to promote cell growth was acquired during carcinoma progression. Schematic, normal mammary gland (**A**), ductal carcinoma in situ (**D**) and invasive carcinoma (**G**). MCF10A, MCF10A-DCIS and MCF10CA1 cells were seeded on plastic, (**B,C,E,F,H,I**) 2mg/ml collagen I (coll I) or (**J**) CAF-CDM for 8 or 6 days under glutamine (Gln) and amino acid (AA) starvation, fixed, stained with DRAQ5 (**B,C,E,F,H,I**) or Hoechst 33342 (**J**) and imaged with a Licor Odyssey system (**B,C,E,F,H,I**) or ImageXpress micro (**J**). Signal intensity was calculated by Image Studio Lite software (**B,C,E,F,H,I**) or MetaXpress software (**J**). Values are mean ± SEM from 3 independent experiments (the black dots in the bar graphs represent the mean of individual experiments). *p<0.05, **** p<0.0001 (**B,H,J**) 2way ANOVA, Tukey’s multiple comparisons test. (**I**) Kruskal-Wallis, Dunn’s multiple comparisons test.

### Collagen I and Matrigel increased cell division under amino acid starvation

The ECM-dependent increase in cell number could be mediated by a stimulation of cell division or an inhibition of cell death, or both. To investigate this, we performed 5-Ethynyl-2’-deoxyuridine (EdU) incorporation experiments, as this thymidine analogue allows the identification of cells which passed through DNA synthesis and mitosis. Cells were starved for 6 days and then incubated with EdU for 2 days (**Figure 3A**). As shown in figure 3, both glutamine and amino acid starvation significantly reduced the percentage of EdU positive cells on plastic, while the presence of collagen I (**Figure 3B**) or Matrigel (**Figure 3C**) resulted in a significant increase in EdU positive cells. Consistent with the proliferation data, the presence of ECM did not affect EdU incorporation in complete media (**Figure 3B,C**). To elucidate whether the ECM also played an anti-apoptotic role, MDA-MB-231 cell death was assessed by measuring the percentage of cells positive for an apoptosis marker, activated caspase-3/7 (**Figure S3A**), after three and eight days of amino acid starvation. Although the apoptosis rate increased between day three and day eight in both complete and amino acid-free media, and amino acid starvation resulted in increased cell death, there was no significant difference between the apoptosis rate on collagen I and Matrigel compared to plastic either at day three (**Figure S3B**) or day eight (**Figure S3C**) in amino acid depleted media. Similar results were obtained in MCF10CA1 cells after three (**Figure S3D**) and six (**Figure S3E**) days of amino acid starvation. Thus, our data indicate that collagen I and Matrigel induced invasive breast cancer cell proliferation under amino acid starvation, without having an anti-apoptotic effect.

**Figure 3.**
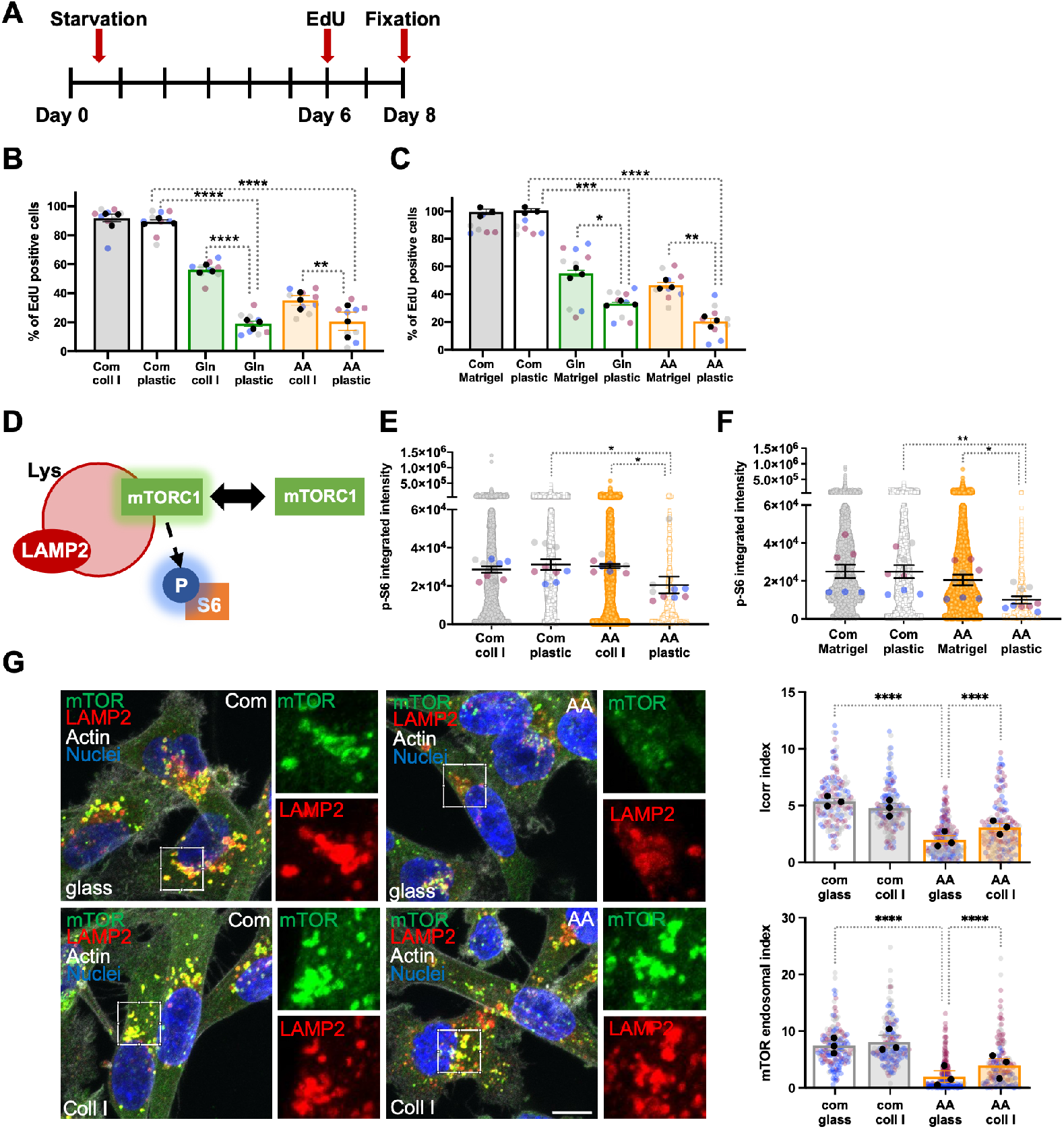
Collagen I and Matrigel induced cell proliferation and rescued mTORC1 activity under amino acid starvation. (**A**) Timeline of EdU incorporation experiments. MDA-MB-231 cells were seeded on plastic, (**B,E**) 2mg/ml collagen I (coll I) or (**C,F**) 3mg/ml Matrigel under complete media (Com), glutamine (Gln) or amino acid (AA) starvation. (**B,C**) Cells were incubated with EdU at day 6 post starvation, fixed and stained with Hoechst 33342 and Click iT EdU imaging kit at day 8. Images were collected by ImageXpress micro and analysed by MetaXpress software. (**D**) Schematic, mTORC1 activation on the lysosomal membrane. (**E,F**) Cells were fixed and stained for p-S6 and nuclei at day 3 post starvation. Images were collected by ImageXpress micro and analysed by Costume Module Editor (CME) software. (**G**) MDA-MB-231 cells were seeded on uncoated or 1mg/ml collagen I coated glass-bottomed dishes for 24hr in complete media, kept in complete (com) or amino acid-free (AA) media for 24hr, fixed and stained for mTOR (green), LAMP2 (red), actin (white) and nuclei (blue). Samples were imaged with a Nikon A1 confocal microscope. mTOR/LAMP2 co-localisation (Icorr index) and mTOR endosomal accumulation were quantified with Image J. Scale bar, 10*μ*m. Values are mean ± SEM from 3 independent experiments (the black dots represent the mean of individual experiments). *p<0.05, **p<0.01, ***p<0.001, **** p<0.0001 Kruskal-Wallis, Dunn’s multiple comparisons test.

### Collagen I and Matrigel rescued mTORC1 activity in starved cells

The mammalian target of Rapamycin (mTOR) signalling pathway is a key regulator of anabolic and catabolic processes. The presence of amino acids, glucose and growth factors activates mTOR complex 1 (mTORC1), which triggers downstream anabolic signalling pathways. However, the lack of amino acids and growth factors deactivates mTORC1 and induces catabolism, resulting in lysosome biogenesis and autophagy^21^. Therefore, we investigated whether higher proliferation rate of cells on collagen I and Matrigel was accompanied by a higher mTORC1 activity in starved MDA-MB-231 cells. To do this, we examined the phosphorylation of the ribosomal subunit S6, a key signalling event downstream of mTORC1 activation (**Figure 3D**). MDA-MB-231 cells were starved for three days either on collagen I, Matrigel or plastic, fixed and stained for phospho-S6 (p-S6). Quantifying the intensity of p-S6, as expected, demonstrated that amino acid starvation resulted in a significant reduction in mTORC1 activity on plastic. Interestingly, the presence of ECM fully rescued mTORC1 activation under amino acid starvation (**Figure 3E,F**). mTOR recruitment to the lysosomal membrane has been shown to be integral to mTORC1 activation, as the mTORC1 complex senses amino acid availability in the lysosomes^22^. In agreement with our p-S6 data, amino acid starvation resulted in a significant reduction of mTOR lysosomal recruitment, quantified as co-localisation with the lysosomal marker LAMP2 and mTOR endosomal accumulation. Interestingly, the presence of collagen I led to a significant increase in mTOR lysosomal targeting under amino acid starvation, without affecting mTOR distribution in complete media (**Figure 3G**). Together, given the fact that amino acid availability in lysosomes can promote mTOR recruitment and activation, we can speculate that ECM endocytosis and degradation might result in higher lysosomal amino acid concentrations, supporting mTORC1 translocation and activation.

### Inhibition of ECM internalisation opposed cell growth under amino acid starvation

It was previously shown that, under starvation, cancer cells relied on scavenging extracellular proteins to maintain their survival and growth^11, 15^. Therefore, we set out to characterise the mechanisms controlling ECM internalisation in breast cancer cells. Several endocytic pathways, including clathrin-dependent, caveolin-dependent endocytosis and macropinocytosis have been previously implicated in the internalisation of different ECM components^23^. We developed an high-throughput imaging method, based on pH-rodo labelled ECM and live cell imaging. pH-rodo is a pH sensitive dye, whose fluorescence is minimal at neutral pH but strongly increases in the acidic environment of late endosomes and lysosomes. It has been extensively used in phagocytosis studies^24, 25^. As a first step, we used pharmacological inhibitors targeting dynamin, a GTPase involved in both clathrin and caveolin-mediated endocytosis (dynasore), lipid-raft mediated endocytosis, which includes caveolin-dependendent internalisation (filipin) and macropinocytosis (EIPA) (**Figure 4A**) and we assessed the internalisation of Matrigel, collagen I and NF-generated CDM by MDA-MB-231 cells. As shown in **Figure 4B-C**, the strongest uptake inhibition of all ECM tested was obtained by EIPA treatment, while dynasore decreased Matrigel and CDM endocytosis, but not collagen I and filipin only opposed Matrigel uptake. To validate these data, we used siRNA-mediated downregulation of key endocytosis regulators and assessed Matrigel uptake. Consistent with the inhibitors’ results, the knock-down of dynamin 2/3, caveolin 1/2 and PAK1, a key macropinocytosis regulator^26^, all resulted in a reduction of Matrigel endocytosis (**Figure S4A**), indicating that, although multiple endocytic pathways are likely responsible for the internalisation of complex ECM, macropinocytosis inhibition had the strongest effect on multiple matrices. Indeed, PAK1 knockdown (**Figure S4D**) strongly impaired CAF-CDM (**Figure 4D**), NF-CDM and collagen I uptake in MDA-MB-231 cells (**Figure S4B,C**). To examine whether invasive breast cancer cells had the ability to internalise ECM components under starvation, we tracked the ECM journey inside the cells in the presence and absence of a lysosomal protease inhibitor (E64d), to prevent lysosomal degradation. E64d has been previously shown to promote the intracellular accumulation of lysosomally-delivered proteins, including internalised collagen IV^27^. The uptake of fluorescently-labelled collagen I (**Figure S4E**) and Matrigel (**Figure S4F**) was monitored in complete or amino acid depleted media. The quantification of collagen I and Matrigel uptake demonstrated that, under all nutrient condition, MDA-MB-231 cells significantly accumulated higher amounts of collagen I and Matrigel inside the cells when lysosomal degradation was inhibited (**Figure S4E,F**), indicating that ECM components undergo lysosomal degradation following internalisation. We then wanted to investigate whether the ECM-dependent growth of cancer cells under nutrient deficiency relied on ECM internalisation. To assess this, collagen I and Matrigel coated plates were treated with 10% glutaraldehyde to chemically crosslink the amine groups of ECM components. Our data confirmed that cross-linking completely opposed collagen I and Matrigel internalisation (**Figure S4G-I**) in both MDA-MB-231 and MCF10CA1 cells. Interestingly, while the growth of MDA-MB-231 cells was not affected by matrix cross-linking in complete media (**Figure S4J**), the lack of collagen I and Matrigel uptake due to cross-linking completely opposed cell growth under amino acid starvation (**Figure S4K,L**) in both MDA-MB-231 and MCF10CA1 cells. High matrix cross-linking has been shown to also oppose extracellular ECM degradation mediated by matrix metalloproteinases (MMPs)^28^, therefore we wanted to determine whether the inhibition of ECM-dependent cell growth by cross-linked matrices was due to a reduction in ECM degradation by MMPs (**Figure S5A**). In agreement with previous reports showing that ECM internalisation was not dependent on MMP activity^23^, our data indicated that MMP inhibition by treatment with the broad spectrum inhibitor GM6001 did not reduce collagen I and Matrigel uptake (**Figure S5B**). Consistent with this, MMP inhibition did not affect cell growth under complete media or amino acid starvation on both plastic and collagen I (**Figure S5C-E**). Since our data indicate that macropinocytosis is the main pathway controlling ECM endocytosis in breast cancer cells, we wanted to investigate whether PAK1-mediated ECM uptake was required for ECM dependent cell growth. Consistent with the matrix cross-linking data, PAK1 downregulation, obtained by either siRNA (**Figure 4E-G**) or CRISPRi sgRNA (**Figure 4G**) strongly reduced cell growth under amino acid starvation when cells were seeded on collagen I or CAF-CDM. In addition, treatment with an ATP-competitive PAK inhibitor (FRAX597, **Figure 4H**) also prevented ECM-dependent cell growth, indicating that PAK1 catalytical activity is required for this. Interestingly, PAK1 inhibition did not affect cell proliferation on plastic, while resulted in a small but significant reduction in cell growth on ECM in the presence of complete media (**Figure S4M-O**). Similarly, PAK pharmacological inhibition and PAK1 downregulation substantially reduced MCF10CA1 cell growth on collagen I under amino acid starvation (**Figure 4I,J**). Altogether, these results demonstrate that macropinocytosis-dependent ECM internalisation is necessary for ECM-dependent breast cancer cell growth under amino acid starvation.

**Figure 4.**
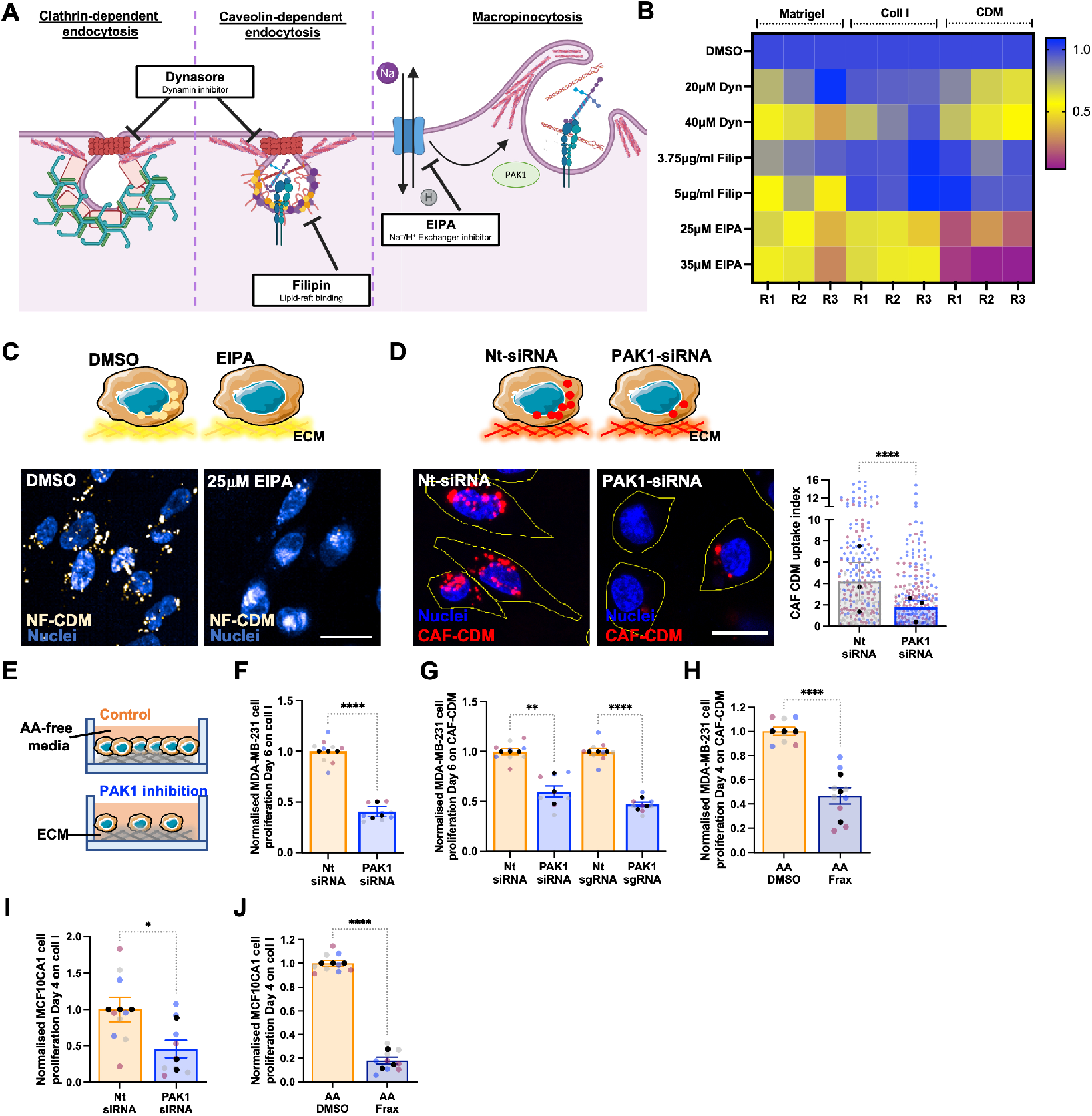
PAK1-mediated ECM macropinocytosis was required for ECM-dependent cell growth under amino acid starvation. (**A**) Schematic, main endocytic pathways and inhibitor targets. (**B,C**) MDA-MB-231 cells were seeded on pH-rodo labelled 0.5mg/ml Matrigel, 0.5mg/ml collagen I (coll I) or NF-CDM (CDM, white) in the presence of DMSO control, 20*μ*M or 40*μ*M dynasore, 3.75*μ*g/ml or 5*μ*g/ml filipin, 25*μ*M or 35*μ*M EIPA for 6hrs, stained with Hoechst 33342 (blue), imaged live with an Opera Phenix microscope and analysed with Columbus software. Scale bar, 20*μ*m. (**D**) MDA-MB-231 cells were transfected with an siRNA targeting PAK1 (PAK1-siRNA) or a non-targeting siRNA control (nt-siRNA), plated on pH-rodo labelled CAF-CDM (red) for 6hr, stained with Hoechst 33342 (blue) and imaged live with a Nikon A1 confocal microscope. Scale bar, 20*μ*m. ECM uptake index was quantified with Image J. (**E**) Schematic, cell proliferation experiments. MDA-MB-231 (**F,G**) and MDA-MB-231 CRSPRi (**G**) cells were plated on 2mg/ml collagen I (coll I, **F**) or CAF-CDM (**G**), transfected with an siRNA targeting PAK1 (PAK1-siRNA), a non-targeting siRNA control (nt-siRNA, **F,G**), a synthetic guide RNA targeting PAK1 (PAK1-sgRNA) or a non-targeting synthetic guide RNA control (nt-sgRNA, **G**) and cultured under amino acid starvation for 6 days. MDA-MB-231 cells (**H**) and MCF10CA1 cells (**J**) were grown on CAF-CDM (**H**) or 2mg/ml collagen I (coll I, **J**) under amino acid (AA) starvation for 4 days in the presence of 3*μ*M (**H**) or 1*μ*M (**J**) FRAX597, fixed and stained with Hoechst 33342. MCF10CA1 cells (**I**) were transfected with an siRNA targeting PAK1 (PAK1-siRNA) or a non-targeting siRNA control (nt-siRNA) and cultured under amino acid (AA) starvation for 4 days. Cells were fixed and stained with Hoechst 33342. Images were collected by ImageXpress micro and analysed by MetaXpress software. Values are mean ± SEM and from 3 independent experiments (the black dots represent the mean of individual experiments). *p<0.05, **p<0.01, **** p<0.0001 Mann-Whitney test (**D,F,H,I,J**) or Kruskal-Wallis, Dunn’s multiple comparisons test (**G**).

### Inhibition of focal adhesion kinase did not affect ECM-dependent cell growth under starvation

ECM cross-linking has been shown to increase matrix stiffness, thereby affecting cell adhesion and integrin signalling^23^. Binding of ECM components to the integrin family of ECM receptors induces the recruitment of focal adhesion (FA) proteins on the integrin intracellular domain, triggering downstream signalling pathways (**Figure 5A**)^29, 30^. Focal adhesion kinase (FAK) is a non-receptor tyrosine kinase located in FAs involved in the regulation of several cell behaviours, including proliferation, migration, and survival. FAK is also over-expressed in some cancers and it has been linked to their aggressiveness^31^. To test whether the inhibition of focal adhesion signalling affected cell growth under nutrient starvation, MDA-MB-231 cells were treated with a FAK inhibitor, PF573228. Firstly, the auto-phosphorylation of FAK Tyr397 was measured in the presence of different PF573228 concentrations, to assess the effectiveness of the inhibitor. Our results revealed that around 80% inhibition of FAK activity was achieved with 0.5*μ*M PF573228 (**Figure 5B,C**). Therefore, to assess the effect of FAK inhibition on cell growth, MD-MB-231 cells were seeded either on collagen I, Matrigel or CAF-generated CDM under complete media or amino acid starvation in the presence of 0.5*μ*M PF573228. Interestingly, we did not observe any significant difference in cell number between the control group and cells treated with the FAK inhibitor, either in complete media or amino acid deprivation (**Figure 5D-F**). In addition, staining for the FA protein paxillin indicated no changes in the quantity and distribution of FAs in cells under amino acid starvation compared to the complete media, on either collagen I or Matrigel, (**Figure 5G,H**) or on cross-linked matrices (**Figure 5I**). Taken together, these data indicate that FAK-dependent signalling events following cell-ECM interaction did not affect cell growth under amino acid starvation and ECM-dependent cell growth was predominantly mediated by ECM internalisation followed by lysosomal degradation.

**Figure 5.**
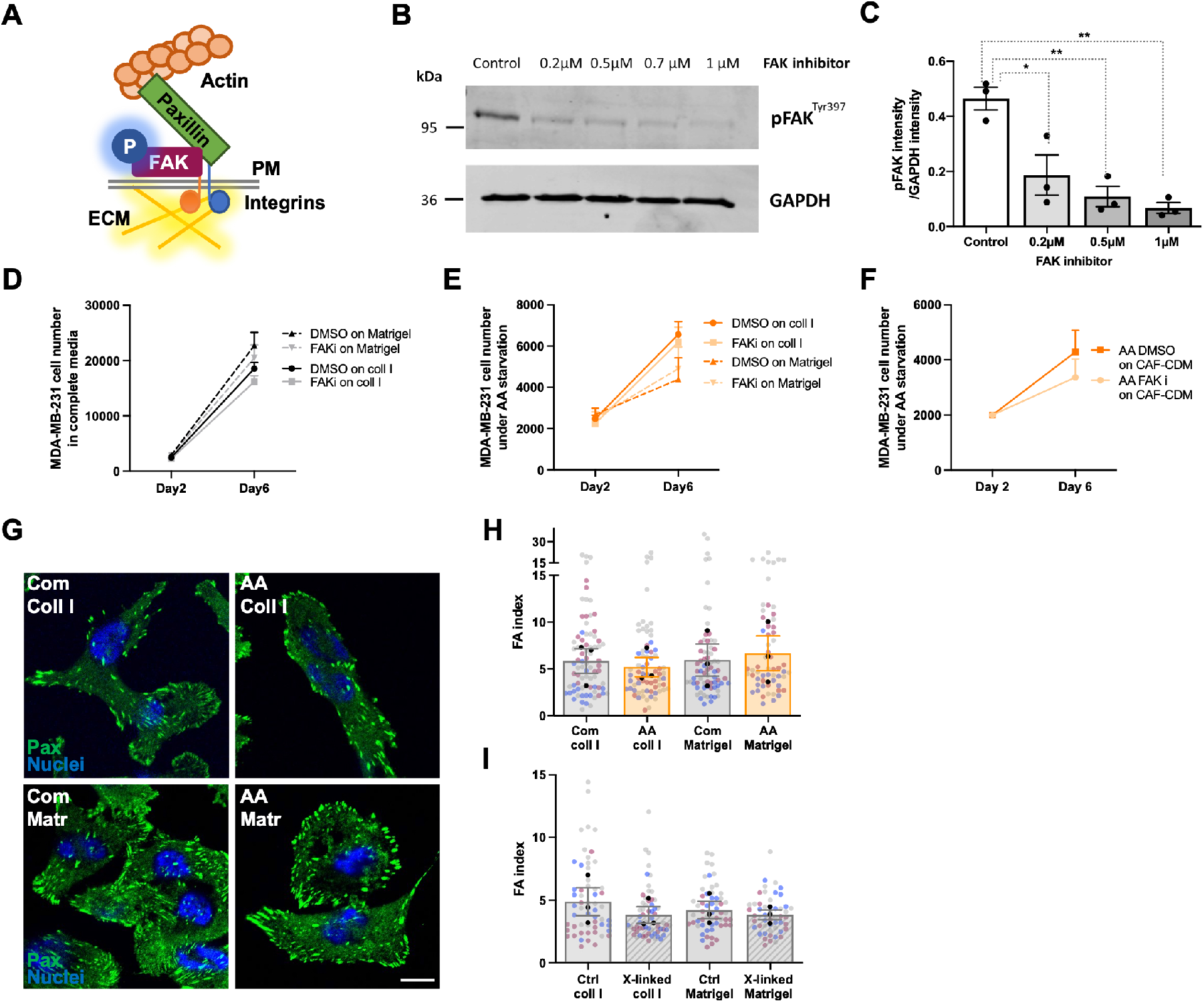
Focal adhesion signalling was not required for ECM-dependent cell growth. (**A**) Schematic, focal adhesions. (**B,C**) MDA-MB-231 cells were suspended in complete media containing either 0.2*μ*M, 0.5*μ*M, 0.7*μ*M or 1*μ*M PF573228 (FAK inhibitor) or DMSO (vehicle) for 30 mins. Cells were seeded on 0.5mg/ml collagen I for 30 mins in the same media, lysed and proteins analysed by Western Blotting. The intensity of the bands was quantified using the Lycor Odyssey system. Values are mean ± SEM from 3 independent experiments. *p<0.05, **p<0.01 Kruskal-Wallis, Dunn’s multiple comparisons test. (**D-F**) MDA-MB-231 cells were seeded on either 2mg/ml collagen I (coll I), 3mg/ml Matrigel, CAF-CDM or plastic under complete media or amino acid starvation for 6 days and treated with 0.5*μ*M PF573228 (FAKi) or DMSO (control) every 2 days. Cells were fixed and stained with Hoechst 33342. Images were collected by ImageXpress micro and analysed by MetaXpress software. Values are mean ± SEM from 3 independent experiments. (**G-I**) MDA-MB-231 cells were plated on 2mg/ml collagen I (coll I) or 3mg/ml Matrigel (Matr), previously cross-linked with 10% glutaraldehyde where indicated (x-linked), for 3 days in complete (Com) or amino acid free (AA) media, fixed and stained for paxillin (green) and nuclei (blue). Bar, 10*μ*m. Images were acquired with a Nikon A1 confocal microscope and quantified with Image J. Values are mean ± SEM from 3 independent experiments (the black dots represent the mean of individual experiments).

### ECM-dependent cell growth was mediated by phenylalanine/tyrosine metabolism under amino acid starvation

It has been previously demonstrated that laminin endocytosis resulted in an increase in intracellular amino acid levels in serum starved epithelial cells^11^, therefore we hypothesised that metabolites derived from ECM internalisation and degradation would result in changes in cellular metabolome. To quantify ECM-dependent differences in the cellular metabolite content, we performed non-targeted direct infusion mass spectrometry of MDA-MB-231 cells either on plastic or CAF-CDM under amino acid starvation for 6 days (**Figure 6A**). On CAF-CDM, we detected 716 upregulated and 749 downregulated metabolites (**Figure 6B**). Metabolomic pathway analysis revealed that phenylalanine and tyrosine metabolism and the pentose phosphate pathway were the most significantly enriched (**Figure 6C**). We then used a similar approach comparing MDA-MB-231 cells grown in plastic, collagen I or Matrigel for 6 days under amino acid starvation (**Figure S6A**). Metabolic pathway analysis revealed that purine metabolism and phenylalanine/tyrosine metabolism were the most upregulated (**Figure S6B-E**). Consistently, when we assessed amino acid intracellular levels, we found that phenylalanine and tyrosine were the most upregulated on all the ECMs tested compared to plastic under amino acid starvation (**Figure 6D**). By performing a metabolic pathway analysis restricted to the metabolites commonly upregulated on all ECMs compared to plastic (**Figure 6E**), tyrosine and phenylalanine metabolism was identified as the most upregulated pathway (**Figure 6F**). Phenylalanine metabolism is composed of a series of metabolic reactions, leading to the production of fumarate, which can enter the tricarboxylic acid (TCA) cycle, leading to energy production (**Figure 6G**). Using targeted ultra-performance liquid chromatography-tandem mass spectrometry in MDA-MB-231 cells grown under amino acid starvation for 6 days, we confirmed that the presence of CAF-CDM and collagen I resulted in increased levels of phenylalanine, tyrosine and fumarate (**Figure 6H,I**). Interestingly, the inhibition of ECM uptake by treatment with the PAK inhibitor FRAX597 resulted in a reduction of phenylalanine and tyrosine in cells grown on collagen I under amino acid starvation, suggesting that ECM endocytosis is required for the upregulation of tyrosine and phenylalanine on ECM under starvation. We reasoned that, if the ECM supported cell growth by promoting phenylalanine catabolism, the knock-down of a central enzyme in this pathway would inhibit ECM-dependent cell growth under amino acid deprivation (**Figure 6K**). To test this, we transiently knocked-down p-hydroxyphenylpyruvate hydroxylase (HPD) and its paralogue p-hydroxyphenylpyruvate hydroxylase-like protein (HPDL) and we assessed cell proliferation on plastic or CAF-CDM, under complete media or amino acid starvation. In both cases, we observed a significant reduction in protein or mRNA expression upon siRNA transfection (**Figure S6O,P**) Strikingly, HPDL knockdown, but not HPD, significantly impaired cell growth under amino acid starvation on CAF-CDM (**Figure 6L**) and on collagen I (**Figure S6G**). Interestingly, both HPD and HPDL knock-down resulted in a small, albeit statistically significant, reduction in cell proliferation on CDM in complete media and opposed cell growth in diluted Plasmax media (**Figure S6F,H**). In addition, HPD down-regulation, but not HPDL, modestly reduced cell numbers on plastic. Consistently CRISPRi-mediated HPDL downregulation opposed cell growth under amino acid starvation on CAF-CDM (**Figure 6M**). Both HPD and HPDL have previously been shown to be involved in multiple metabolic pathways^32, 33^. To assess whether HPDL was controlling phenylalanine catabolism is our settings, we measured intracellular fumarate levels and we showed that HPDL knock-down resulted in a significant reduction in fumarate concentration (**Figure S6J**). To confirm that fumarate could be generate by tyrosine catabolism, we performed an isotope labelling experiment by incubating MDA-MB-231 cells with C13-tyrosine. Interestingly, we detected a higher proportion of C13-labelled fumarate on plastic than on collagen I (**Figure S6K**), suggesting that tyrosine derived from collagen I degradation could contribute to fumarate production, therefore increasing the M+0 pool in cells plated on collagen I. To confirm that HPD/HPDL catalytic activity was required for ECM-dependent cell growth, we treated MDA-MB-231 cells with nitisinone, an FDA-approved 4-hydroxyphenylpyruvate dioxygenase inhibitor currently used to treat hereditary tyrosinemia type 1, and observed a significant reduction in cell numbers on CAF-CDM under amino acid starvation (**Figure 6N**). Consistent with these data, metabolomics analysis identified tyrosine and phenylalanine metabolism amongst the upregulated pathways in MCF10CA1 cells grown on collagen I under amino acid starvation (**Figure S6L,M**). Targeted ultra-performance liquid chromatography-tandem mass spectrometry confirmed increased phenylalanine, tyrosine and fumarate levels in MCF10CA1 cells on collagen I under amino acid starvation, compared to plastic (**Figure S6N**) and HPDL downregulation resulted in a significant inhibition of cell proliferation (**Figure S6Q**). Given the requirement for HPDL to control ECM-dependent cell growth under amino acid starvation, we looked at the relationship between HPDL expression and disease outcome in breast cancer. High expression of HPDL was associated with significantly decreased overall survival (**Figure 6O**). Altogether, these data indicate that HPDL has a key role in controlling ECM-dependent cell growth under amino acid starvation. 3D spheroids have been shown to display significant metabolic changes compared to their 2D counterparts^34^, therefore we assessed whether the ECM was also able to sustain cell proliferation in 3D culture conditions. We first measured the ability of MDA-MB-231 cells grown as spheroids to internalise ECM components, by labelling the collagen I and geltrex mixture with pH-rodo. Live cell imaging demonstrated a significant increase in ECM endocytosis 24hr and 48hr after amino acid starvation, suggesting that nutrient deprivation promotes ECM uptake in 3D systems (**Figure 7A,B**). Morevover, we investigated whether tyrosine catabolism was required for the growth of MDA-MB-231 cell spheroids under amino acid starvation. Consistent with our 2D experiments, we observed that amino acid starved spheroids retain substantial EdU incorporation, while the presence of the HPD/HPDL inhibitor nitisinone resulted in a statistically significant reduction in the percentage of EdU positive cells, without affecting the invasive abilities of the cells (**Figure 7C**). Together with our 2D results, these data demonstrate that tyrosine catabolism plays a central role in promoting ECM-dependent cancer cell proliferation, in both 2D and 3D environments.

**Figure 6.**
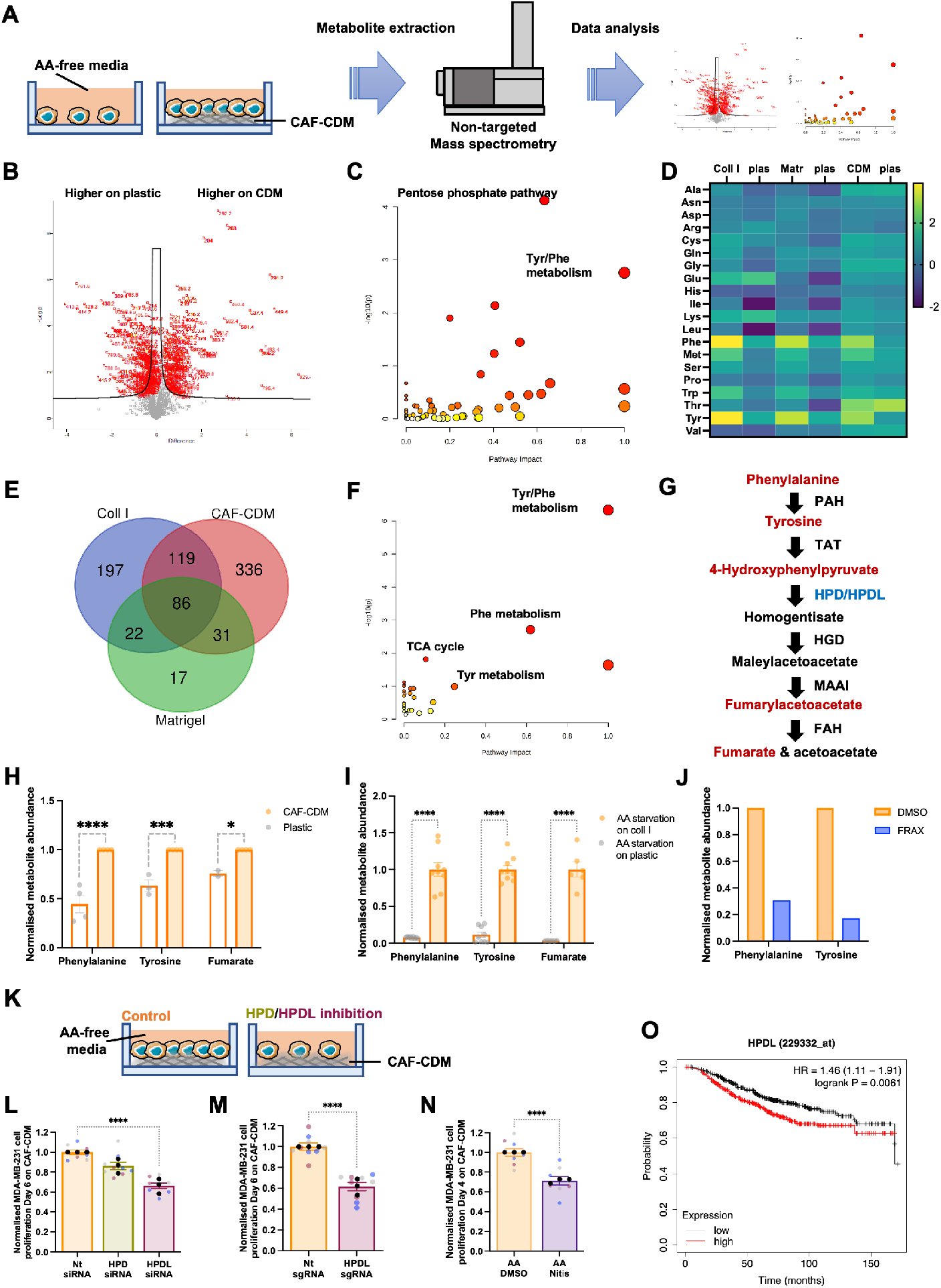
ECM-dependent cell growth was mediated by tyrosine catabolism. (**A**) Metabolomics workflow. MDA-MB-231 cells were plated on plastic or CAF-CDM for 6 days in amino acid-free media. Metabolites were extracted and quantified by non-targeted mass spectrometry. Volcano plot (**B**), enriched metabolic pathways (**C**) and changes in amino acid levels (**D**) are presented. (**E**) Venny diagram and metabolic pathway analysis (**F**) of the statistically significant upregulated metabolites in cells plated on 2mg/ml collagen I (coll I), CAF-CDM or 3mg/ml Matrigel under amino acid starvation for 6 days. (**G**) Schematic, phenylalanine catabolism. (**H-J**) Metabolites were prepared as in (A), or in the presence of 2*μ*M FRAX597, and the levels of phenylalanine, tyrosine and fumarate were measured by targeted mass spectrometry. (**K,L**) MDA-MB-231 cells were plated on CAF-CDM, transfected with siRNA targeting HPD (HPD siRNA), siRNA targeting HPDL (HPDL siRNA) or non-targeting siRNA control (Nt siRNA) and cultured under amino acid (AA) starvation for 6 days. Cells were fixed and stained with Hoechst 33342. Images were collected by ImageXpress micro and analysed by MetaXpress software. (**M**) MDA-MB-231 CRSPRi cells were plated on CAF-CDM, transfected with a synthetic guide RNA targeting HPDL (HPDL sgRNA) and a non-targeting synthetic guide RNA control (Nt sgRNA) and analysed as in (L). (**N**) MDA-MB-231 cells were grown on CAF-CDM under amino acid (AA) starvation for 4 days in the presence of 40*μ*M Nitisinone (Nitis), and analysed as in (L). *p<0.05, *** p<0.001, **** p<0.0001 2way ANOVA, Tukey’s multiple comparisons test (**H,I**), Mann-Whitney test (**N,M**) or Kruskal-Wallis, Dunn’s multiple comparisons test (**L**). (**O**) 943 Breast cancer patients were stratified into low and high HPDL expressors based on median gene expression. The Kaplan-Meier analysis compared the survival outcome of patients with tumours expressing high levels of HPDL (red), with those expressing low HPDL levels (black).

**Figure 7.**
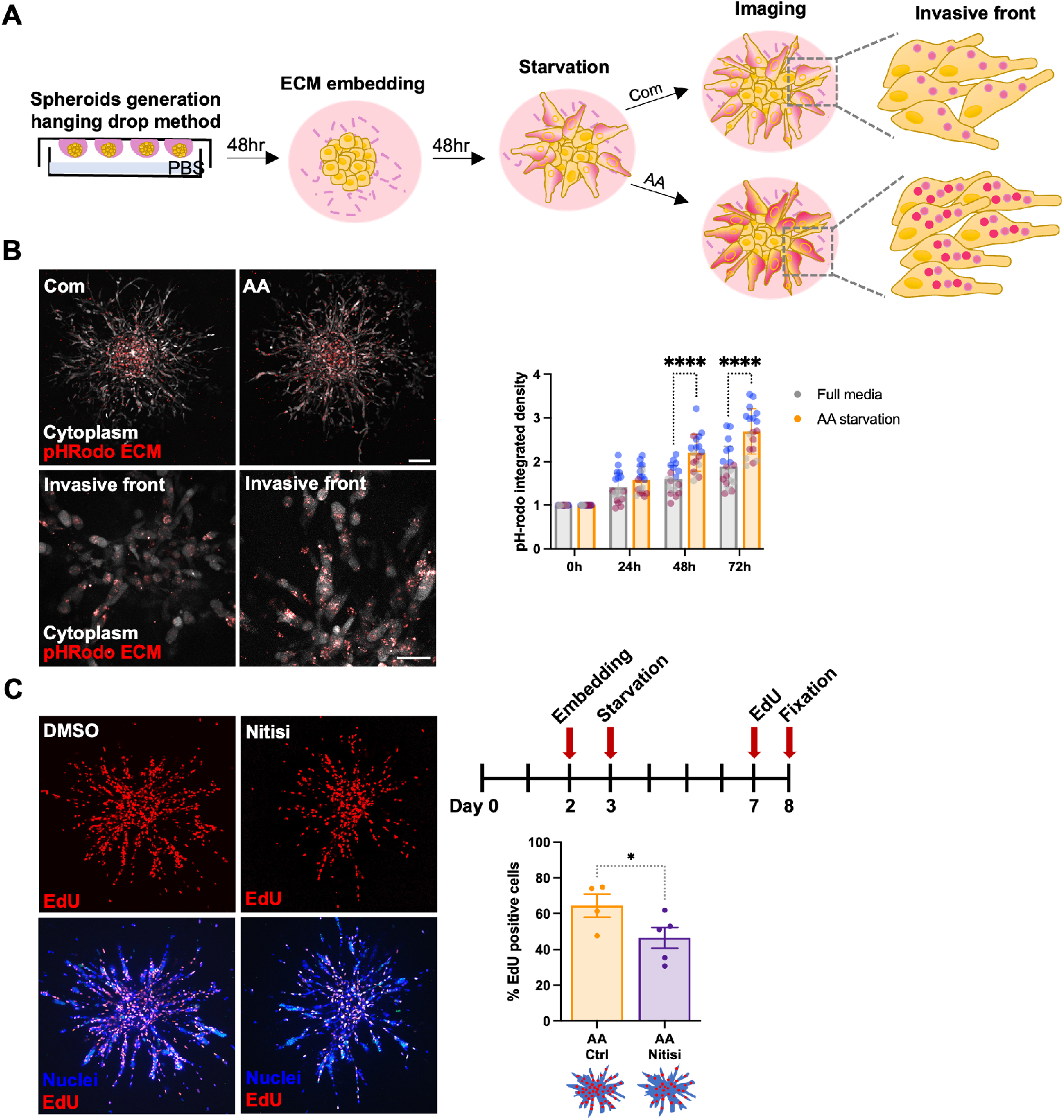
Tyrosine catabolism supported cell growth in 3D systems. (**A**) Schematic, ECM uptake in 3D spheroids. (**B**) MDA-MB-231 cell spheroids were generated by the hanging drop method, embedded in 3mg/ml pH-rodo labelled collagen I and geltrex (50:50) mixture for 2 days, starved of amino acid (AA) or kept in complete media (Com) for 2 days and imaged live with a Nikon A1 confocal microscope. Representative images of whole spheroids and higher magnification of the invasive front are shown. Scale bar, 132*μ*m and 42*μ*m, respectively. Values are mean ± SD from 3 independent experiments, **** p<0.0001 2way ANOVA, Tukey’s multiple comparisons test. (**C**) MDA-MB-231 cell spheroids were generated as in (B) in unlabelled 3mg/ml collagen I and geltrex (50:50) mixture, starved in amino acid-free (AA) media for 5 days, incubated with EdU for 1 day, fixed and stained with Hoechst 33342 and Click iT EdU imaging kit at day 6. The percentage of EdU-positive cells was measured with Image J. Values are mean ± SEM from 3 independent experiments * p=0.0286 Mann-Whitney test.

### ECM-derived tyrosine catabolism supports pancreatic cancer cell growth

Collagens have previously been shown to sustain PDAC cell growth under glucose and glutamine starvation, by stimulating proline metabolism^15^. To assess whether the presence of the ECM was able to promote the proliferation of PDAC cells under amino acid starvation, we monitored the growth of PANC1 cells plated on plastic or on collagen I. In agreement with our results in breast cancer cells, cell proliferation was significantly increased in the presence of collagen I (**Figure 8A**), while no difference in growth was detected in complete media regardless of the substrate the cells were seeded on (**Figure S7A**). In addition, collagen I was internalised and lysosomally degraded in both complete and amino acid-free media, as in the presence of the lysosomal inhibitor E64d we detected a significantly higher amount of internalised collagen I compared to the DMSO control (**Figure S7B**). Furthermore, inhibition of ECM macropinocytosis by PAK1 inhibition significantly opposed collagen I uptake (**Figure 8B**) and PANC1 cells growth on collagen I under both complete media and amino acid starvation (**Figure 8C,S7C,D**), without having any effect on cell proliferation on plastic (**Figure S7C**). These data indicate that ECM internalisation supports PDAC cell growth both in complete media and under amino acid deprivation. Metabolomics analysis identified several metabolites upregulated on collagen I under amino starvation, compared to plastic (**Figure 8D,E**). Metabolomic pathway analysis highlighted tyrosine and phenylalanine metabolism among the enriched pathways (**Figure 8F**) and targeted ultra-performance liquid chromatography-tandem mass spectrometry confirmed increased levels of phenylalanine, tyrosine and fumarate in PANC1 cells grown on collagen I compared to plastic under amino acid starvation (**Figure 8G**). Consistently, the blockade of tyrosine catabolism by HPD and HPDL knock-down significantly impaired cell growth on collagen I in both complete media and amino acid deprivation, while no effect was observed on plastic (**Figure 8H,I, S7E**). Looking at the relationship between HPDL expression and disease outcome in PDAC, we found that high expression of HPDL was associated with significantly decreased overall survival (**Figure 8J**). Altogether, these data indicate that HPDL-mediated tyrosine catabolism plays an important role in supporting ECM-dependent cell growth also in PDAC cells.

**Figure 8.**
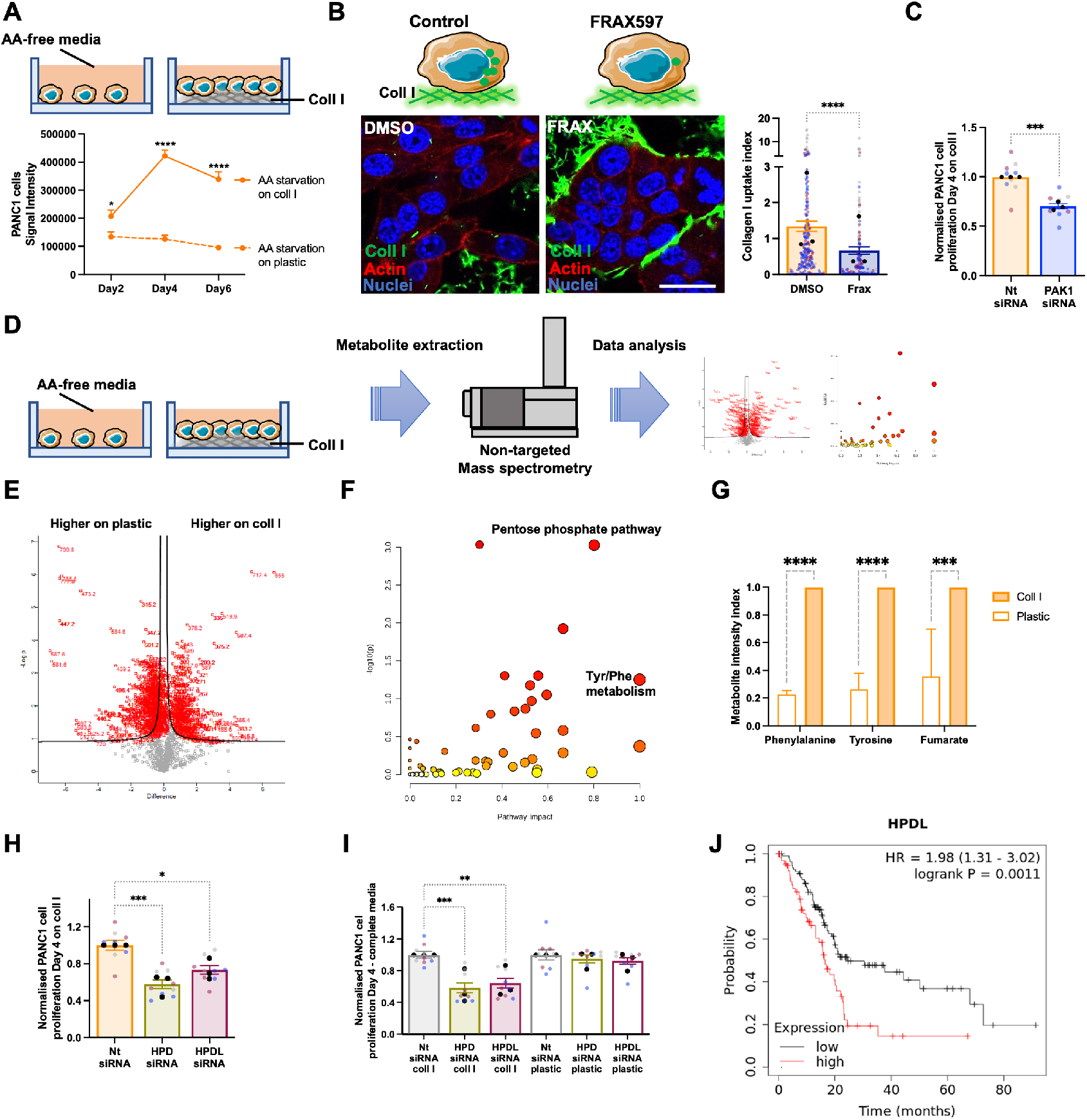
ECM-derived tyrosine catabolism supports pancreatic cancer cell growth. (**A**) PANC1 cells were seeded on plastic or 2mg/ml collagen I (coll I) for 6 days under amino acid (AA) starvation, fixed, stained with DRAQ5 and imaged with a Licor Odyssey system. Signal intensity was calculated by Image Studio Lite software. (**B**) PANC1 cells were plated under amino acid depleted (AA) media on NHS-fluorescein labelled 2mg/ml collagen I (coll I) coated dishes for 3 days, in the presence of the lysosomal inhibitor E64d (20*μ*M) and 2*μ*M FRAX597. Cells were fixed and stained for actin (red) and nuclei (blue). Samples were imaged with a Nikon A1 confocal microscope and ECM uptake index was calculated with image J. Bar, 20*μ*m. About 200 cells per condition in three independent experiments were analysed, the black dots represent the mean of individual experiments. **** p<0.0001 Kruskal-Wallis, Dunn’s multiple comparisons test. (**B**) PANC1 cells were plated on 2mg/ml collagen I (coll I), transfected with an siRNA targeting PAK1 (PAK1-siRNA) or a non-targeting siRNA control (nt-siRNA) and cultured under amino acid starvation for 4 days. Cells were fixed and stained with Hoechst 33342. Images were collected by ImageXpress micro and analysed by MetaXpress software. Values are mean ± SEM and from three independent experiments (the black dots represent the mean of individual experiments). ***p<0.001 Mann-Whitney test. (**D**) Metabolomics workflow. PANC1 cells were plated on plastic or collagen I for 6 days in amino acid-free media. Metabolites were extracted and quantified by non-targeted mass spectrometry. Volcano plot (**E**) and enriched metabolic pathways (**F**) are presented. (**G**) Metabolites were prepared as in (D) and the levels of phenylalanine, tyrosine and fumarate were measured by targeted mass spectrometry. PANC1 cells were plated on 2mg/ml collagen I (coll I, **H,I**) or plastic (**I**), transfected with an siRNA targeting HPD (HPD-siRNA), an siRNA targeting HPDL (HPDL-siRNA) or a non-targeting siRNA control (nt-siRNA), cultured under amino acid starvation (**H**) or complete media (**I**) for 4 days and analysed as in (C). *p<0.05, **p<0.01, ***p<0.001 Kruskal-Wallis, Dunn’s multiple comparisons test. (**J**) 177 Pancreatic ductal adenocarcinoma patients were stratified into low and high HPDL expressors based on median gene expression. The Kaplan-Meier analysis compared the survival outcome of patients with tumours expressing high levels of HPDL (red), with those expressing low HPDL levels (black).

## Discussion

Breast and pancreatic cancer cells are embedded into a highly fibrotic microenvironment enriched in several ECM components^5^. Different studies suggested that cancer cells rely on extracellular protein internalisation during starvation^9, 11–14^. However, to date no studies investigated the behaviour of invasive breast cancer cells in response to the presence of ECM under nutrient starvation. Here, we demonstrated that ECM internalisation and lysosomal degradation provide a source of amino acids, which supports cell growth in a phenylalanine and tyrosine metabolism-dependent manner, when invasive breast and pancreatic cancer cells experience amino acid starvation (**Figure 9**). Importantly, we showed that the ability to use ECM to sustain cell growth under starvation was progressively acquired during carcinoma progression, as the ECM was unable to rescue cell growth in non-transformed MCF10A; while Matrigel, but not collagen I, promoted the growth of non-invasive MCF10A-DCIS cells under glutamine, but not amino acid, deprivation. Non-invasive breast cancer cells reside inside the mammary duct and are in contact with the basement membrane, but not with stromal collagen I^35^. Since Matrigel is a basement membrane extract, our data suggest that MCF10A-DCIS cells are only able to take advantage of basement membrane components to support their growth under starvation. Consistent with our data in MDA-MB-231, highly invasive and metastatic MCF10CA1 cell growth was supported by the ECM under both starvation conditions. In contrast, it was previously demonstrated that the growth of serum-starved MCF10A was promoted by the incubation with soluble laminin^11^, suggesting that the role of the ECM in supporting non-transformed epithelial cell growth could be dependent on the type of starvation experienced by the cells. Indeed, we observed a partial rescue of MCF10A cell growth in the presence of Matrigel under serum starvation (data not shown).

**Figure 9.**
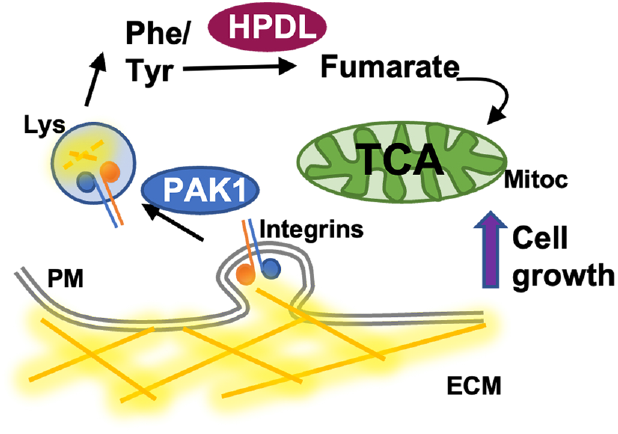
Working model. Invasive breast and pancreatic cancer cells internalised ECM components through PAK1-mediated macropinocytosis, resulting in ECM lysosomal degradation. This upregulated phenylalanine/tyrosine catabolism, leading to an HPDL-dependent increased production of fumarate, supporting cell proliferation under amino acid starvation. ECM, extracellular matrix; PM, plasma membrane; Lys, lysosome; Mitoc, mitochondria.

It has recently been reported that amino acid starvation induced MT1-MMP-dependent ECM degradation, through the inhibition of MT1-MMP endocytosis^36^. Here, we showed that MMP inhibition did not oppose ECM internalisation and ECM-dependent cell growth. However, Colombero et al. assessed ECM degradation within the first 24hr after amino acid starvation^36^, while we focused on later time points (3 and 6 days). Therefore, it is possible that the induction of ECM degradation is an early response to amino acid starvation, which is not required to sustain cell proliferation at later time points. In addition, collagen I endocytosis was recently shown not to be affected by GM6001 treatment in lung adenocarcinoma cells^37^.

We showed that the presence of ECM fully rescued mTORC1 activation under amino acid starvation. This is in agreement with several previous studies showing that mTORC1 could be reactivated by extracellular proteins under amino acid deficiency^11, 12^ through a positive feedback loop, as inhibition of mTORC1 induced extracellular protein scavenging^11, 12^. Similarly, we have previously demonstrated that mTORC1 inhibition promoted fibronectin internalisation in ovarian cancer cells^13^. Furthermore, our data showed that the presence of ECM elevated mTORC1 activity in starved cells to the same level as cells in complete media, whereas, the ECM only partially rescued cell growth under starvation, indicating that growth is still limited despite a complete activation of mTORC1. It is important to note that we used only one readout of mTORC1 activation (S6 phosphorylation), therefore we cannot exclude the possibility that the ECM was unable to rescue other signalling pathways downstream of mTORC1. Interestingly, it has been previously shown that elevated mTORC1 activity restrains growth if cells rely on scavenging extracellular proteins as a source of amino acids. This is because high mTORC1 activity results in higher translation rate and cells relying on extracellular proteins cannot support the high rate of both protein translation and growth at the same time, due to their reduced access to free amino acids. Therefore, mTORC1 reactivation resulted in lower growth in cells using extracellular proteins as amino acid source^38^. Previous literature showed that, in pancreatic cancer cells, collagen-dependent cell growth under glucose and glutamine starvation was coupled with activation of ERK, but not mTORC1, signalling^15^, suggesting that the ECM supports cell growth via triggering distinct downstream signalling pathways in different cancer types.

We characterised the endocityc pathways controlling the internalisation of different ECM components and, consistently with previous observation^23, 37^, while different pathways contributed to ECM uptake, macropinocytosis played the most important role in controlling the internalisation of different ECM types. PAK1 is a well-established regulator of macropinocytosis^26^ and it has been shown to be upregulated in a variety of cancer types, including breast cancer^39, 40^. PAK1 can be activated downstream of multiple signalling pathways, including the small GTPases Rac and Cdc42 as well as Phosphatydil inositol 3 kinase (PI3K)/AKT. By using both matrix cross-linking and PAK1 inhibition, we demonstrated that ECM internalisation is required for ECM-dependent cell growth under amino acid starvation. Interestingly, PI3K, but not Rac or Cdc42, seems to be required for ECM internalisation and ECM-dependent cell growth under amino acid starvation (data not shown).

Phenylalanine and tyrosine metabolism has been shown to be implicated several cancer types. Interestingly, a comprehensive metabolomic study identified the upregulation of phenylalanine and tyrosine catabolism in invasive ductal carcinoma compared to benign tumours and normal mammary tissues^41^. HPD has been shown to be upregulated in invasive breast cancer and, to a lesser extent, DCIS compared to normal tissues and its expression correlated with poor prognosis^42^. In lung cancer, high HPD expression correlated with poor prognosis. HPD has been reported to promote cell growth in vitro and in vivo, by controlling the flux through the pentose phosphate pathway, in a tyrosine metabolism-independent manner^32^. It is therefore possible that HPD controls cell growth on plastic via a similar mechanism in our system as well. In colon adenocarcinoma cells, it has been shown that HPDL knock-down reduced oxygen consumption, implying that it could play a role in controlling mitochondria functions. While we could detect a reduction in fumarate levels upon HPDL knock-down, in colon cancer cells HPDL has been shown to localise to mitochondria and its knock-down did not result in impaired tyrosine catabolism^33^, suggesting that, similarly to HPD, HPDL could be involved in multiple metabolic pathways. Interestingly, breast cancer patient survival analysis indicate that high expression of HPDL correlated with a reduced overall survival. Future work will characterise the mechanism through which HPDL controls ECM-dependent cell proliferation. Our data suggest that ECM-derived fumarate, generated through tyrosine catabolism, fuels the TCA cycle, as we could detect the accumulation of other TCA intermediates, including α-ketoglutarate, succinate and malate in cells growing on ECM under amino acid starvation (data not shown). In agreement with this, it has been previously reported that, in the presence of collagen I, a significant proportion of the TCA intermediate citrate is not labelled by glucose or glutamine, suggesting that the collagen matrix could represent a fuel source in the TCA cycle^43^. However, fumarate has also been proposed to behave as an oncometabolite, and its accumulation has been shown to promote the stabilisation of hypoxia inducible factor (HIF)^44^. Further work will elucidate the molecular mechanisms through which the increase in fumarate levels generated by the activation of tyrosine catabolism supports ECM-dependent cell growth under amino acid starvation.

Furthermore, we showed that the ECM was similarly able to support PDAC cell growth under amino acid starvation, through a mechanism involving PAK1-mediated ECM uptake and tyrosine catabolism. Interestingly, in the context of PDAC, HPD and HPDL were required for cell growth both in complete media and under starvation, suggesting a more prominent and nutrient level-independent role of these enzymes in controlling cell proliferation in this cancer type. While in vitro both HPD and HPDL down-regulation opposed ECM-dependent cell growth on collagen I, only the expression of HPDL, but not HPD, correlated with PDAC patient prognosis.

In conclusion, our findings demonstrated that, under amino acid starvation, invasive breast and pancreatic cancer cells, but not non-transformed mammary epithelial cells and non-invasive DCIS cells, internalised and degraded ECM components. This resulted in an increase in tyrosine catabolism, leading to the production of fumarate. Therefore, HPDL-mediated tyrosine catabolism could represent a metabolic vulnerability of cancer cells thriving in a nutrient-deprived microenvironment.

## Methods

### Reagents

Primary antibodies for Phospho-S6 Ribosomal Protein ser235/236 and GAPDH were from Cell Signalling, Phospho-FAK (Tyr397), HPD and HPDL from Thermo Fisher, *β* 1 integrin (clone TS2/16) from BioLegend, Paxilin from BD-bioscience and PAK1 from Proteintech. Secondary antibodies Alexa-fluor 594 anti-Rabbit IgG, Alexa-fluor 488 anti-Rabbit IgG, Alexa-fluor 594 anti-Mouse IgG and Alexa-fluor 488 anti-Mouse IgG were from Cell Signalling, IRDye® 800CW and IRDye® 680CW were from LI-COR. Alexa fluor TM 555 Phalloidin, Click-iT EdU Imaging Kits, CellEvent™ Caspase-3/7 Green Detection kit, NHS-Fluorescein, NHS-Alexa Fluor 555, pH-rodo iFL STP ester red and Hoechst 33342 were from Invitrogen. DRAQ5 was from LI-COR. Collagen I and Matrigel were from Corning. All media and dialyzed FBS were from Gibco, except for DMEM with no amino acid which was from US Biological life science and Plasmax which was kindly provided by Dr Tardito, CRUK Beatson Institute, Glasgow. The details of Plasmax composition were previously described by Vande Voorde et al 2019^19^. E64d (Aloxistatin) and PF573228 were from AdooQ Bioscience. GM6001 was from APEXBIO. Dynasore, Filipin and EIPA were from Sigma. FRAX 597 was from bio-Techne. Nitisinone was from MCE. C13-tyrosine was from Sigma-Aldrich.

### Cell culture

MDA-MB-231 cells, PANC1 cells, Telomerase immortalised normal fibroblasts (NFs) and cancer associated fibroblasts (CAFs), generated in Professor Akira Orimo’s lab, Paterson Institute, Manchester, were cultured in High glucose Dulbecco’s Modified Eagle’s Medium (DMEM) supplemented with 10% fetal bovine serum (FBS) and 1% penicillin streptomycin (PS). MCF10A, MCF10A-DCIS and MCF10CA1 were a kind gift from Prof Giorgio Scita, IFOM, Milan (Italy). MCF10A-DCIS cells were cultured in DMEM/F12 supplemented with 5% Horse serum (HS), 20 ng/ml epithelial growth factor (EGF) and 1% PS. MCF10A cells were cultured in DMEM/F12 supplemented with 5% HS, 20 ng/ml EGF, 0.5mg/ml hydrocortisone, 10*μ*g/ml insulin and 1% PS. MCF10CA1 cells were cultured in DMEM/F12 supplemented with 2.5% HS, 20 ng/ml EGF, 0.2mg/ml hydrocortisone, 10*μ*g/ml insulin and 1% PS. Cells were grown at 5% CO_2_ and 37°C and passaged every 3 to 4 days.

### ECM preparation

To coat plates/dishes, collagen I and Matrigel were diluted with cold PBS on ice to 2mg/ml and 3mg/ml concentrations respectively and incubated for 3 hrs at 5% CO_2_, and 37°C for polymerization. Cell derived matrix (CDM) was generated from NFs and CAFs as described in^16^. Briefly, plates were coated with 0.2% gelatin in PBS and incubated at 37°C for 1 hr and cross-linked with 1% glutaraldehyde in PBS at room temperature (RT) for 30 mins. Glutaraldehyde was removed and quenched in 1M glycine at RT for 20 mins, followed by two PBS washes, and an incubation in complete media at 5% CO_2_ and 37°C for 30 mins. CAFs or NFs were seeded and incubated at 5% CO_2_, 37°C until fully confluent. The media was replaced with full growth media containing 50*μ*g/ml ascorbic acid, and it was refreshed every two days. NFs were kept in media supplemented with ascorbic acid for six days while CAFs for seven days. The media was aspirated, and the cells washed once with PBS containing calcium and magnesium. The cells were incubated with the extraction buffer containing 20mM NH_4_OH and 0.5% Triton X-100 in PBS containing calcium and magnesium for 2 mins at RT until no intact cells were visible by phase contrast microscopy. Residual DNA was digested with 10*μ*g/ml DNase I at 5% CO_2_, 37°C for 1 hr. The CDMs were kept in PBS containing calcium and magnesium at 4°C.

### ECM cross-linking

The ECMs were prepared as described above and treated with 10% glutaraldehyde for 30 mins at RT, followed by two PBS washes. The cross-linker was quenched in 1M glycine at RT for 20 mins, followed by two PBS washes. Cross-linked ECMs were kept in PBS at 5% CO_2_ and 37°C over-night.

### Starvation conditions

The media used for the starvations were supplemented with either 10% dialyzed FBS (DFBS, for MDA-MB-231 cells); 10% HS, 10*μ*g/ml insulin and 20ng/ml EGF (for MCF10A cells); 5% HS and 20ng/ml EGF (for MCF10A-DCIS cells) or 2.5% HS, 20 ng/ml EGF, 0.2mg/ml hydrocortisone and 10*μ*g/ml insulin (for MCF10CA1 cells). Cells were plated in complete media for 5 hrs, then the media was replaced with the starvation media as indicated in table 1.

**Table 1.**
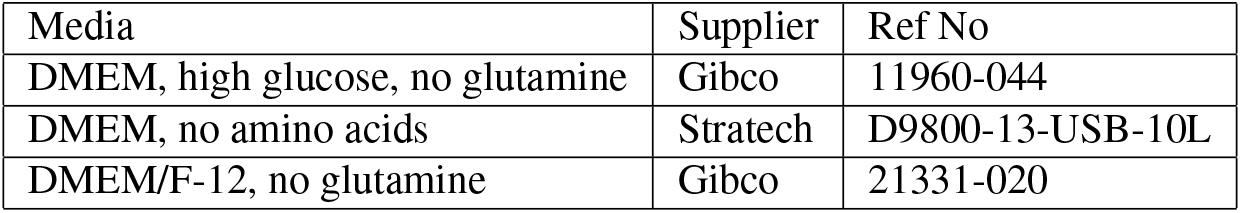
Starvation media.

### Proliferation assays

96-well plates were coated with 15*μ*l/well of 2mg/ml collagen I or 3mg/ml Matrigel. Three wells of 96-well plate were allocated to each condition as technical replicates. 10^3^ cells/well (MDA-MB-231, MCF10A, MCF10A-DCIS cells) or 400 cells/well (MCF10CA1) were seeded in complete growth media. After a 5 hr incubation in 5% CO_2_ and 37°C, the media was replaced with 200*μ*l of the starvation media, in the presence of 10*μ*M GM6001, 0.5*μ*M PF573228, 3*μ*M FRAX597, 40*μ*M Nitisinone or DMSO control where indicated. The inhibitors were supplemented every two days, up to day 4 or 6. Cells were fixed by adding 4% paraformaldehyde (PFA) for 15 mins at RT followed by two PBS washes. Cells were stained with either DRAQ5 (assays on collagen I and Matrigel) or Hoechst 33342 (assays on CDMs, where the thickness and structure of the matrices resulted in a very high background and accurate readings could not be obtained with DRAQ5). 5*μ*M DRAQ5 in PBS was added to the cells for 1 hr at RT with gentle rocking. Cells were washed twice with PBS for 30 mins to avoid background fluorescence and kept in PBS for imaging. DRAQ5 was detected by the 700nm channel with 200*μ*m resolution with an Odyssey Sa instrument. The signal intensity (total intensity minus total background) of each well was quantified with Image Studio Lite software. Alternatively, cells were fixed with 4% PFA containing 10*μ*g/ml Hoechst 33342 for 15 mins, followed by two PBS washes. Images were acquired by ImageXpress micro with a 2x objective and the whole area of each well was covered. The images were analysed with MetaXpress and Costume Module Editor software (CME) in the Sheffield RNAi Screening Facility (SRSF). CAF/NF-CDM was generated in 96-well plates and cells were seeded as described above. Cells were fixed with 4% PFA containing 10*μ*g/ml Hoechst 33342 for 15 mins, washed twice with PBS, permeabilized with 0.25% Triton X-100 for 5 mins and stained with Phalloidin Alexa Fluor 488 (1:400 in PBS) for 30 mins. Cells were washed twice with PBS and were left in PBS for imaging. Images were collected by ImageXpress micro and analysed by MetaXpress and CME software.

### EdU incorporation essays

96-well plates were coated with ECM and 10^3^ MDA-MB-231 cells/well were seeded in full growth media. After a 5 hr incubation at 5% CO_2_ and 37°C, the full growth media was replaced with 200*μ*l of the starvation media. At day six post starvation, cells were incubated with 5*μ*M EdU for 2 days at 5% CO_2_ and 37°C, fixed with 4% PFA containing 10*μ*g/ml Hoechst 33342 for 15 mins at RT and permeabilised with 0.25% Triton X-100 for 5 mins. Cells were incubated with EdU detection cocktail (Invitrogen, Click-iT EdU Alexa Fluor 555) for 30 min at RT with gentle rocking. Cells were washed twice with PBS and were kept in PBS for imaging. Images were collected by ImageXpress micro with a 2x objective and quantified with MetaXpress and CME software.

### Caspase-3/7 detection

96-well plates were coated with ECM and 10^3^ MDA-MB-231 or MCF10CA1 cells/well were seeded in full growth media. After a 5 hr incubation at 5% CO_2_ and 37°C, the full growth media was replaced with 200*μ*L of the starvation media. Cells were kept under starvation for 3 or 6/8 days, then the media were replaced by 5*μ* M CellEvent™ Caspase-3/7 Green Detection Reagent diluted in PBS, containing calcium and magnesium, and 5% DFBS for 1.5 hrs at 5% CO_2_ and 37°C. Cells were fixed with 4% PFA containing 10*μ*g/ml Hoechst 33342 for 15 mins at RT, washed and left in PBS for imaging. Images were collected by ImageXpress micro with a 10x objective and analysed with MetaXpress and CME software.

### Immunofluorescence

10^3^ MDA-MB-231 cells/well were seeded in full growth media in 96-well plates were coated with ECM. After a 5 hr incubation at 5% CO_2_ and 37°C, the full growth media was replaced with 200*μ*L of the starvation media. After 3 days, cells were fixed with 4% PFA containing 10*μ*g/ml Hoechst 33342 for 15 mins at RT and permeabilised with 0.25% Triton X-100 for 5 mins. Unspecific binding sites were blocked by incubating with 3% BSA at RT with gentle rocking for 1 hr. Cells were incubated with anti-Phospho-S6 Ribosomal protein antibody (1:100 in PBS) overnight at 4°C. The cells were washed twice with PBS for 5 to 10 mins and incubated with secondary antibody (anti-Rabbit IgG Alexa Fluor 594, 1:1000 in PBS) for 1hr at RT with gentle rocking. Cells were washed twice with PBS for 10 to 30 mins and were kept in PBS for imaging. Images were collected by ImageXpress micro with a 10x objective and analysed with MetaXpress and CME software.

4×10^4^ MDA-MB-231 cells were seeded on ECM coated 3.5cm^2^ glass-bottomed dishes and incubated at 5% CO_2_ and 37°C for 5 or 24 hrs. The full growth media was then replaced with the starvation media. After 1 or 3 days, cells were fixed with 4% PFA for 15 mins and permeabilised with 0.25% Triton-×100 for 10mins, followed by 1hr incubation in 3% BSA (blocking buffer). Cells were incubated with anti-Paxillin (1:200 in PBS), anti-HPDL (1:100 in blocking buffer), anti-mTOR (1:100 in blocking buffer) or anti-LAMP2 (1:100 in blocking buffer) primary antibodies at RT for 1 hr. Cells were washed three times with PBS for 20 mins and incubated with Alexa-Fluor 488 anti-Mouse IgG, Alexa-Fluor 594 anti-Mouse IgG or Alexa-Fluor 488 anti-rabbirt IgG secondary antibodies (1:1000 in blocking buffer) for 1 hr at RT with gentle shaking and protected from light. 2-3 drops of Vectashield mounting medium containing DAPI were added to the dishes, which were then sealed with parafilm and kept at 4°C. Cells were visualised by confocal microscopy using a Nikon A1 microscope and 60x 1.4NA oil immersion objective. The focal adhesion index and mTOR endosomal index were calculated dividing the area covered by the paxillin or mTOR staining by the total cell area. mTOR/LAMP2 co-localisation (Icorr index) was calculated using the co-localisation analysis plug in developed by^45^.

### ECM uptake

3.5cm^2^ glass-bottomed dishes were coated with 1mg/ml or 2mg/ml collagen I or 3mg/ml Matrigel and incubated at 5% CO_2_, and 37°C for 3 hrs. ECMs were cross-linked where indicated and labelled with 10*μ*g/ml Fluorescein or Alexa Fluor 555 succinimidyl esters (NHS) for 1 hr at RT on a gentle rocker. 10^5^ MDA-MB-231 cells per dish were seeded in full growth media. After a 5 hr incubation in 5% CO_2_ and 37°C, full growth media were replaced with 1ml of the indicated starvation media in the presence of 20*μ*M E64d or DMSO. E64d and DMSO were added after two days and cells were fixed at day three by adding 4% PFA for 15 mins at RT. Cells were permeabilised with 0.25% Triton X-100 for 5 mins and incubated with Phalloidin Alexa Fluor 555 (1:400 in PBS) for 10 mins. 2-3 drops of Vectashield mounting medium containing DAPI were added to the dishes, which were then sealed with parafilm and kept at 4°C. Cells were visualised by confocal microscopy using a Nikon A1 microscope and 60x 1.4NA oil immersion objective. For the high-throughput imaging of ECM endocytosis, 10^4^ MDA-MB-231 cells/well were seeded on pH-rodo labelled 0.5mg/ml Matrigel, 0.5mg/ml collagen I or TIF-CDM in 384-well plates in the presence of DMSO control, 20*μ*M or 40*μ*M Dynasore, 3.75*μ*g/ml or 5*μ*g/ml filipin, 25*μ*M or 35*μ*M EIPA for 6 hrs, labelled with 10*μ*g/ml Hoechst 33342 and imaged live with a 63x water immersion objective with an Opera Phenix microscope. 2.5*μ*l of 500nM siGENOME siRNA smart pool and 2.5*μ*l Opti-MEM per well was added into CellCarrier Ultra 384 well plates (Perkin Elmer). 4.95*μ*l Opti-MEM was incubated with 0.05*μ*l Dharmafect IV for 5min. 5*μ*l of the Dharmafect IV solution was added into each well. Plates were incubated for 20min on gentle rocker at RT. 3×10^3^ cells were seeded in 40*μ*l DMEM containing 10% FBS. The final concentration of the siRNA was 25nM. Cells were kept at 5% CO_2_ and 37°C for 72 hr. Transfected cells were transferred into pH-rodo-labelled 0.5mg/ml Matrigel-coated CellCarrier Ultra 384 well plates. Cells were incubated for 6hr, stained for Hoechst 33342 and imaged live with a 40x water immersion objective with Opera Phenix microscope. Images were analysed with Columbus software. ECM uptake was quantified with Fiji^46^ as described in^47^.

### Western Blotting

MDA-MB-231 cells were trypsinized and 10^6^ cells were collected in each Falcon tube. Cells were treated with 0.2, 0.5, 0.7 and 1*μ*M P573228 for 30 mins at 5% CO_2_ and 37°C, while they were kept in suspension. Cells were transferred to a 6-well plate which had been coated with 900*μ*L of 0.5mg/ml collagen I. Cells were allowed to adhere for 30 mins at 5% CO_2_ and 37°C. The media was aspirated, and cells were put on ice. After two washes with ice-cold PBS, 100*μ*l per well of lysis buffer (50mM Tris pH 7 and 1% SDS) were added. Lysates were collected and transferred to QiaShredder columns, which were spun for 5 mins at 6000rpm. The filter was discarded, and the extracted proteins were loaded into 12% acrylamide gels and run at 100V for 2.5 hrs. Proteins were transferred to FL-PVDF membranes in transfer buffer for 75 mins at 100V. Membranes were washed three times with TBS-T (Tris-Buffered Saline, 0.1% Tween) and blocked in 5% w/v skimmed milk powder in TBS-T for 1 hr at RT. Membranes were then incubated with primary antibodies, 1:500 GAPDH and 1:1000 pFAK in TBS-T with 5% w/v skimmed milk overnight at 4°C. To validate the knock-down efficiency, lysates were run on 4-15% Mini Protean TGX stain Free Protein gels. Membranes were incubated with 1:1000 GAPDH, α-tubulin, PAK1 and HPD primary antibodies in TBS-T with 5%w/v skimmed milk overnight at 4°C. Membranes were incubated with secondary antibodies, IRDye® 800CW anti-mouse IgG for GAPDH and α-tubulin (1:30,000) and IRDye® 680CW anti-rabbit IgG for pFAK, PAK1 and HPD (1:20,000) in TBS-T with 0.01% SDS, for 1 hr at RT on the rocker. Membranes were washed three times in TBS-T for 10 min on the rocker at RT followed by being rinsed with water. Images were taken by a Licor Odyssey Sa system. Band intensity was quantified with Image Studio Lite software.

### Non-targeted metabolite profiling

MDA-MB-231, MCF10CA1 and PANC1 cells were seeded on 96- or 6-well plates coated with ECM under full growth media. After a 5 hr incubation at 5% CO_2_ and 37°C, the full growth media was replaced with the starvation media. After 6 days, the media was removed, and cells were washed with ice-cold PBS three times. The PBS were aspirated very carefully after the last wash to remove every remaining drop of PBS. The extraction solution (cold; 5 MeOH: 3 AcN: 2 H_2_O) was added for 5 mins at 4°C with low agitation. Metabolites were transferred to eppendorf tubes and centrifuged at 4°C at 14000 rpm for 10 mins. Samples were transferred into HPLC vials and directly injected into a Waters G2 Synapt mass spectrometer in electrospray mode within the Sheffield Faculty of Science Biological Mass Spectrometry Facility. The samples were run in positive and negative modes. Three technical replicates of each sample were run. The dataset only includes peaks present in all three replicates. The data is binned into 0.2amu m/z bins and the m/z in each bin is used to identify putative IDs using the HumanCyc database. The mass spectrometry data were analysed using Perseus software version 1.5.6.0. Metabolic pathway analysis was performed in MetaboAnalyst 5.0 (https://www.metaboanalyst.ca/), p<0.05.

### Targeted metabolomics

MDA-MB-231, MCF10CA1 and PANC1 cells were seeded on 6-well plates coated with ECM in full growth media. After a 5 hr incubation at 5% CO_2_ and 37°C, full growth media was replaced with amino acid-free media. Cells were kept under starvation for 6 days, the media was removed and metabolites were extracted as described above. Then, a mass spectrometer Waters Synapt G2-Si coupled to Waters Acquity UPLC was used to separate phenylalanine, tyrosine and fumarate with Column Waters BEH Amide 150×2.1mm. The injection was 10*μ*l. The flow rate was 0.4ml/min. The samples were run in negative mode. To identify the targeted metabolites, the retention time and the mass of the compound were matched with the standards.

### Isotope labelling

6-well plates were coated with 2mg/ml collagen I or left uncoated. 7×10^4^ MDA-MB-231 cells/well were seeded in complete growth media. After a 5 hr incubation in 5% CO_2_ and 37°C, the media was replaced with 4ml of amino acid depleted media, in the presence of 0.4mM C13-tyrosine for 6 days. The metabolites were extracted and analysed as described above by targeted mass spectrometry.

### Transfection

Glass-bottomed 96-well plates were coated with 15*μ*l 2mg/ml collagen I and 20*μ*l of 150nM Dharmacon ON TARGET plus siRNA smart pool or Dharmacon CRISPRmod CRISPRi (inhibition) synthetic sgRNA (**Table 2**) were added on each well. 0.24*μ*l of Dharmafect 1 (DF1) was diluted in 19.76*μ*l of DMEM. 20*μ*l of diluted DF1 were transferred to each well. Plates were incubated for 30 mins at RT. 3×10^3^ cells in 60*μ*l DMEM media containing 10% FBS but no antibiotics were seeded in each well. The final concentration of siRNA was 30nM. Cells were kept at 5% CO_2_ and 37°C over-night. The full growth media was replaced by 200*μ*l of fresh complete or amino acid-depleted media for up to 6 days. At day 2 and 6, cells were fixed and incubated with 10*μ*g/ml Hoechst 33342. Images were collected by ImageXpress micro and analysed by MetaXpress and CME software. 10*μ*l 5*μ*M siRNA or sgRNA were mixed with 190*μ*l Opti-MEM into each well of a 6-well plate. 198*μ*l Opti-MEM and 2*μ*l Dharmafect I were mixed and incubated for 5min at RT. 200*μ*l of the Opti-MEM Dharmafect I mix were added on top of the siRNA and incubated for 20min on a rocker. 4×10^5^ cells in 1.6ml were added into each well. Cells were incubated at 37°C and 5% CO_2_ for 72 hr, either lysed for western blotting or seeded onto pH-rodo-labelled CDMs for 6 hr and imaged live with a Nikon A1 confocal microscope, 60x 1.4NA oil immersion objective.

**Table 2.** siRNAs and sgRNAs.

| Type | Target | Ref No |
| --- | --- | --- |
| ON-TARGETplus siRNA smartpool | PAK1 | L-003521-00-0005 |
| si-GENOME siRNA smartpool | PAK1 | M-003521-04-0005 |
| CRISPRmod CRISPRi sgRNA | PAK1 | CF-003521-01-0005 |
| ON-TARGETplus siRNA smartpool | HPD | L-019648-00-0005 |
| ON-TARGETplus siRNA smartpool | HPDL | L-014985-01-0005 |
| CRISPRmod CRISPRi sgRNA | HPDL | CF-014985-01-0005 |
| ON-TARGETplus siRNA | Non-targeting siRNA 4 | D-001810-04-20 |
| siGENOME siRNA | Non-targeting siRNA 5 | D-001210-05-05 |
| CRISPRmod CRISPRi sgRNA | Non-targeting smartpool | U-009550-10-05 |
| siGENOME siRNA smartpool | Clathrin heavy chain (CHC17) | M-004001-00-0005 |
| siGENOME siRNA smartpool | Dynamin 1 | M-003940-01-0005 |
| siGENOME siRNA smartpool | Dynamin 2 | M-004007-03-0005 |
| siGENOME siRNA smartpool | Dynamin 3 | M-013931-00-0005 |
| siGENOME siRNA smartpool | Caveolin 1 | M-003467-01-0005 |
| siGENOME siRNA smartpool | Caveolin 2 | M-010958-00-0005 |
| siGENOME siRNA smartpool | Flotillin 1 | M-010636-00-0005 |
| siGENOME siRNA smartpool | Flotillin 2 | M-003666-01-0005 |
| siGENOME siRNA smartpool | GRAF 1 | M-008426-01-005 |

### MDA-MB-231 CRISPRi cell generation

3×10^5^ cells/well were seeded into a 12-well plate. Confluent cells were transduced with CRISPRmod CRISPRi dCas9-SALL1-SDS3 lentivirus particles, which expression is under the hCMV promoter (cat VCAS10124), as per protocol (https://horizondiscovery.com/-/media/Files/Horizon/resources/Technical-manuals/edit-r-lentiviral-sgrna-manual.pdf). Cells were incubated at 37°C for 6 hr with 250*μ*l of serum free media containing CRISPRi lentivirus at a multiplicity of infection (MOI) of 0.3. Afterwards, 750*μ*l complete media was added on top of the cells. After 48 hr, cells were collected and a serial dilution was performed. Clones were grown for 2 weeks in the presence of 15*μ*g/ml blasticidin. 12 clones were isolated with Scienceware® cloning discs (Z374431-100EA) and allowed to grow. Clone 8 was picked for future assays based on the efficiency of sgRNA-mediated protein downregulation.

### 3D spheroids

To generate MDA-MB-231 cells stably expressing GFP, 8×10^5^ cells/well were seeded into a 6-well plate in 2ml of complete media without antibiotics. Confluent cells were transfected with 2.5*μ*g of pSBtet-GB GFP Luciferase plasmid and 0.25*μ*g of the sleeping beauty transposon plasmid, pCMV(CAT)T7-SB100. 250*μ*l of OptiMEM, 5*μ*l p3000 and 3.75*μ*l Lipofectamine 3000, together with both plasmids was added on top of the 2ml. Media was changed after 6 hr. After 48 hr, cells were selected with 3*μ*g/ml blasticidin for 2 weeks and FACS sorted.

3D spheroids were generated by the hanging drop method. 500 cells (for proliferation assay) or 1000 cells (for 3D uptake) per 20*μ*l drop containing 4.8mg/ml methylcellulose (Sigma-Aldrich) and 20*μ*g/ml soluble collagen I (BioEngineering) were pipetted on the lid of tissue culture dishes. Lids were turned and put on top of the bottom reservoir of the dish, which was filled with PBS to prevent evaporation. After 48 hr, spheroids were embedded in 40*μ*l of 3mg/ml rat tail collagen I (Corning) and 3mg/ml geltrex (Gibco). For 3D uptake assays, 1/5 of the matrix solution was labelled with 20*μ*g/ml pHrodo containing 0.1M NaHCO_3_. Spheroids were incubated at 37°C for 20min and then media was added into the dishes. For 3D uptake assays, media was changed to 10% DFBS DMEM and 10% DFBS amino acid free DMEM after 44 hr of embedding. Cells were imaged live every 24 hr until 72 hr post-embedding. 3D uptake was assessed by quantifying the pH-rodo-ECM integrated intensity per spheroid, data was normalized to day 0. For 3D proliferation assay, media was changed to 10% DFBS DMEM and 10% DFBS amino acid free DMEM the day after embedding. For drug treatments, 40*μ*M Nitisinone and the corresponding vehicle DMSO was added. On day 5 post-embedding, spheroids were incubated with 10*μ*M EdU for 1 day at 37°C and 5% CO_2_. At day 6, spheroids were fixed by adding 4%PFA containing 10*μ*g/ml Hoechst 33342 for 20 mins at 37°C followed by two PBS washes. They were permeabilized for 2hrs at RT by IF wash containing 0.2% Triton-X 100, 0.04% Tween-20 and 0.1%BSA in PBS followed by two PBS washes. Spheroids were incubated with EdU detection cocktail (Invitrogen, Click-iT EdU Alexa Fluor 555) over-night at 4°C. They were washed twice with PBS and were kept in PBS for imaging with a Nikon A1 confocal microscope.

### Survival analysis

The survival analysis was performed using Kaplan-Meier plotter (https://kmplot.com/analysis/)^48^, which can assess the effect of genes of interest on survival in 21 cancer types, including breast and pancreatic cancer. Sources for the databases include Gene Expression Omnibus (GEO), European Genome-Phenome Archive (EGA), and The Cancer Genome Atlas (TCGA).

### Statistical analysis

Graphs were created by GraphPad Prism software (version 9) as super-plots^49^, where single data points from individual experiment are presented in the same shade (grey, maroon and blue) and the mean is represented by black dots. To compare more than two data set, one-way ANOVA (Kruskal-Wallis, Dunn’s multiple comparisons test) was used when there was one independent variable. Two-way ANOVA (Tukey’s multiple comparisons test) was performed when there were two independent variables. Statistical analysis for non-targeted metabolic profiling was performed by Perseus software and used Student t-test (SO = 0.1 and false discovery rate (FDR) = 0.05).

## Abbreviation list

BM: basement membrane
CAF: cancer-associated fibroblast
CDM: cell-derived matrix
DCIS: ductal carcinoma in situ
ECM: extracellular matrix
EdU: 5-Ethynyl-2’-deoxyuridine
FA: focal adhesion
FAK: focal adhesion kinase
HPD: p-hydroxyphenylpyruvate hydroxylase
HPDL: p-hydroxyphenylpyruvate hydroxylase-like protein
MMP: matrix metalloproteinase
mTOR: mammalial target of Rapamycin
mTORC1: mammalial target of Rapamycin complex 1
NF: normal fibroblast
TCA: tricarboxylic acid cycle
TME: tumour microenvironment.

## Acknowledgements

Metabolomics was performed at the Faculty of Science Biological Mass Spectrometry Facility, University of Sheffield (biOMICS). Imaging work was performed at the Wolfson Light Microscopy Facility, University of Sheffield, using the Nikon A1 confocal/TIRF microscope. High-throughput imaging was performed in the Sheffield RNAi Screening Facility and in the RNAi screening facility in IMCB, Singapore. qPCR analysis was performed in collaboration with the Tsakiridis lab at the University of Sheffield. We would like to acknowledge Marga Albu for developing the Image J macro used for ECM uptake quantification and the ‘ECM and stroma’ group at the University of Sheffield for providing valuable feedback which helped shaping this study. M.N., B.Y. and E.R. are funded by CRUK (C52879/A29144). M.L.M. is funded by Sheffield/ARAP PhD program. The Wolfson Light Microscopy Facility, University of Sheffield, is funded by the Wellcome Trust (grant WT093134AIA).

## Author contributions statement

M.N. and E.R. conceived and performed experiments; B.Y. performed the MCF10CA1 experiments; M.L.M. designed and performed the high-throuput screen for ECM uptake, PAK1 KD ECM uptake, the 3D ECM uptake and generated the stable cell lines used in this study; H.W. performed the metabolomics experiments and analysed the results; FB supervised the high-throughput imaging. All authors reviewed the manuscript.

