## Supplementary figures for "The extracellular matrix supports cancer cell growth under amino acid starvation by promoting tyrosine catabolism"

### Additional information

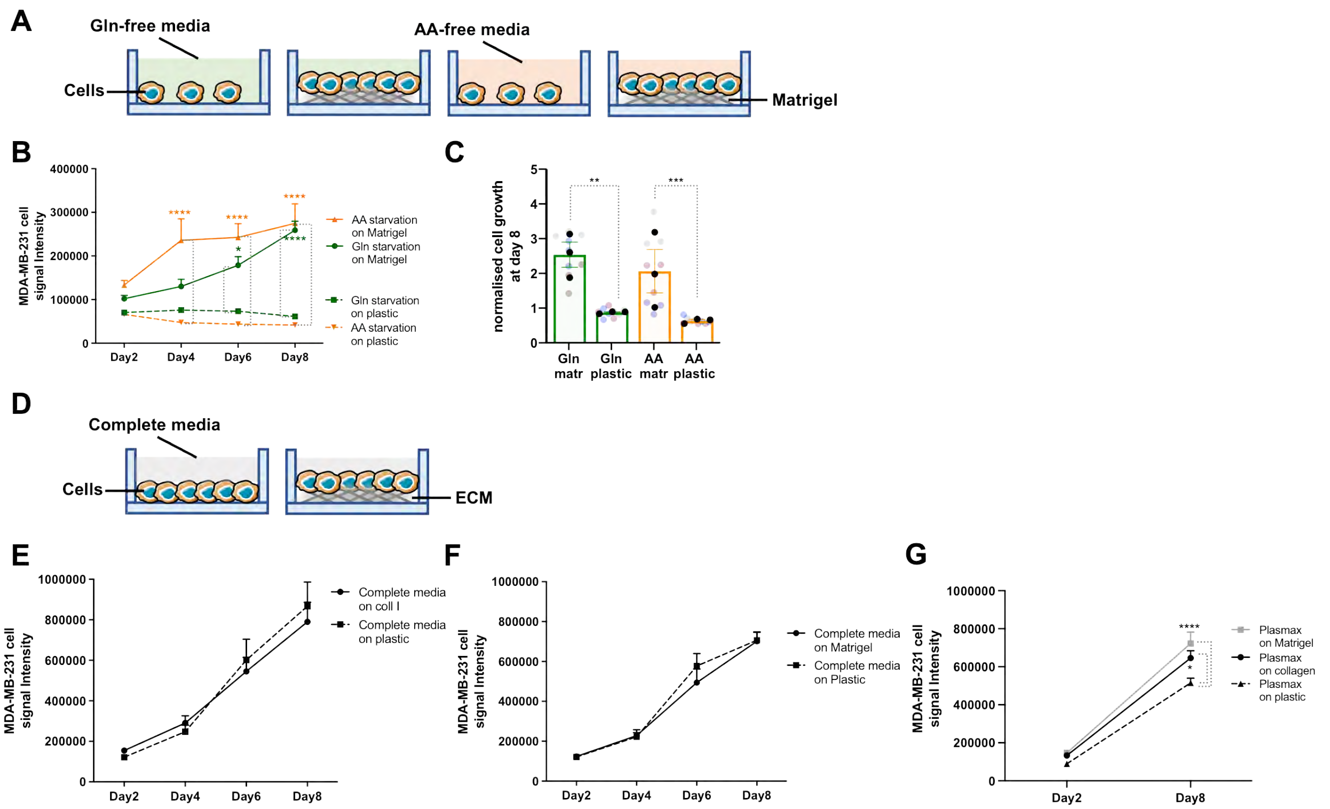


**Figure S1. The ECM had limited effect on cell growth in complete media**. (**A,D**) Schematic, cell proliferation experiments. (**B,C**) MDA-MB-231 cells were seeded on plastic or 3mg/ml Matrigel (matr) for 8 days under glutamine (Gln) or amino acid (AA) starvation, fixed, stained with DRAQ5 and imaged with a Licor Odyssey system. MDA-MB-231 cells were seeded on plastic, (**E**) 2mg/ml collagen I (coll I) or (**F**) 3mg/ml Matrigel for 8 days in complete media or (**G**) in Plasmax media, fixed, stained with DRAQ5 and imaged with a Licor Odyssey system. Signal intensity was calculated by Image Studio Lite software. Values are mean ± SEM and are representative of three independent experiments (the black dots in the bar graphs represent the mean of individual experiments). *p<0.05, **p<0.01, ***p< 0.001, **** p<0.0001 (**B,G**) 2way ANOVA, Tukey's multiple comparisons test. (**C**) Kruskal-Wallis, Dunn's multiple comparisons test.

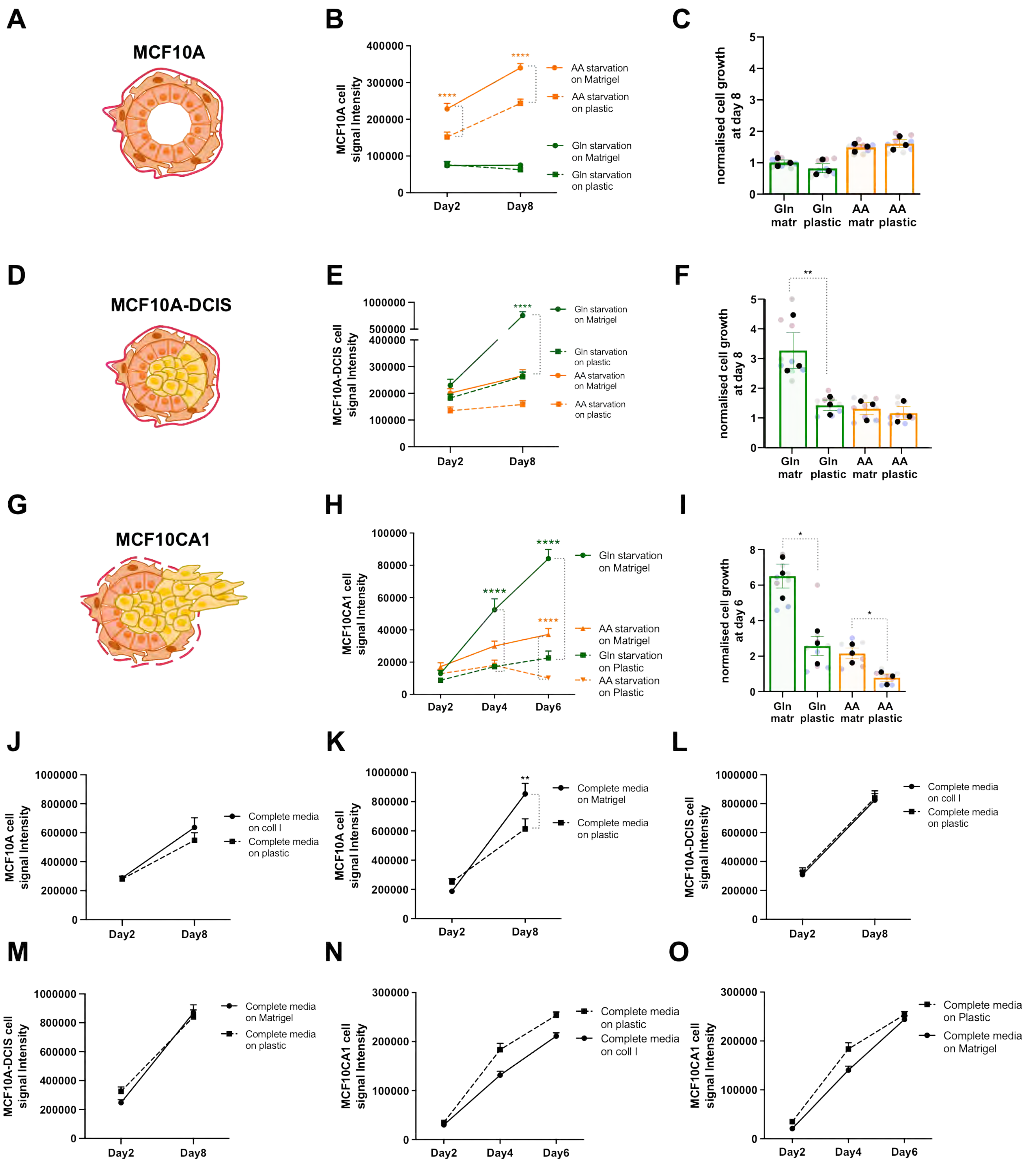

**Figure S2. The ECM had limited effect on cell growth in complete media**. MCF10A (**A-C,J,K**), MCF10DCIS (**D-F,L,M**) and MCF10CA1 (**G-I,N,O**) cells were seeded on plastic, 2mg/ml collagen I (coll I) or 3mg/ml Matrigel for 6 or 8 days under glutamine (Gln) or amino acid (AA) starvation (**A-I**), or in complete media (**J-O**), fixed, stained with DRAQ5 and imaged with a Licor Odyssey system. Signal intensity was calculated by Image Studio Lite software. Values are mean ± SEM and are representative of three independent experiments (the black dots in the bar graphs represent the mean of individual experiments). *p<0.05, **p<0.01, **** p<0.0001 (**B,E,H,K**) 2way ANOVA, Tukey's multiple comparisons test.(**F,I**) Kruskal-Wallis, Dunn's multiple comparisons test.

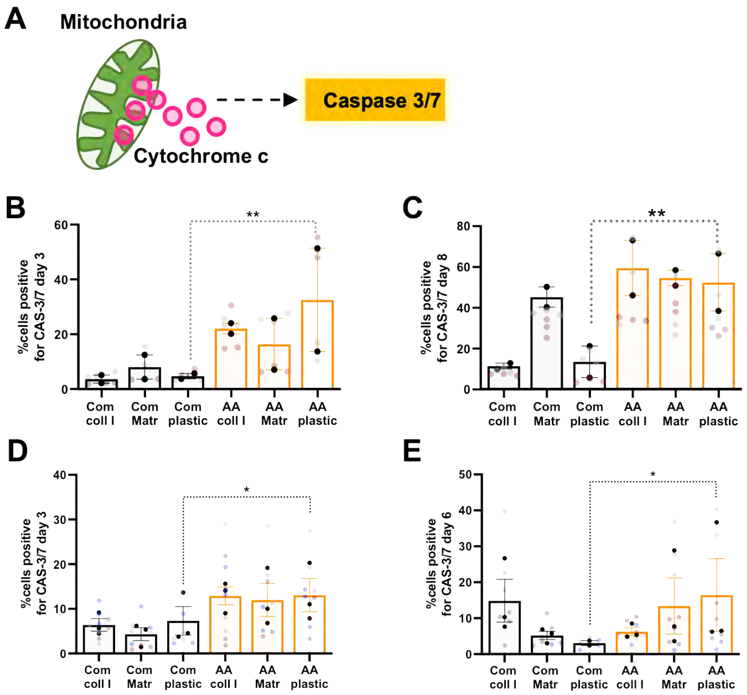


**Figure S3. The ECM did not affect apoptosis under starvation**. (**A**) Schematic, caspase3/7 activation. MDA-MB-231 cells were seeded on plastic, 2mg/ml collagen I (coll I) or 3mg/ml Matrigel (Matr) for (**B**) 3 or (**C**) 8 days in complete media (Com) or amino acid-free media (AA). MCF10CA1 cells were seeded on plastic, 2mg/ml collagen I (coll I) or 3mg/ml Matrigel (Matr) for (**D**) 3 or (**E**) 6 days in complete media (Com) or amino acid-free media (AA). Cells were fixed and stained for activated caspase-3/7 (CAS-3/7). Images were collected by ImageXpress micro and analysed by CME software. Values are mean ±SEM of at least 6 replicates from at least 2 independent experiments (the black dots represent the mean of individual experiments). *p<0.05, **p<0.01 Kruskal-Wallis, Dunn's multiple comparisons test.

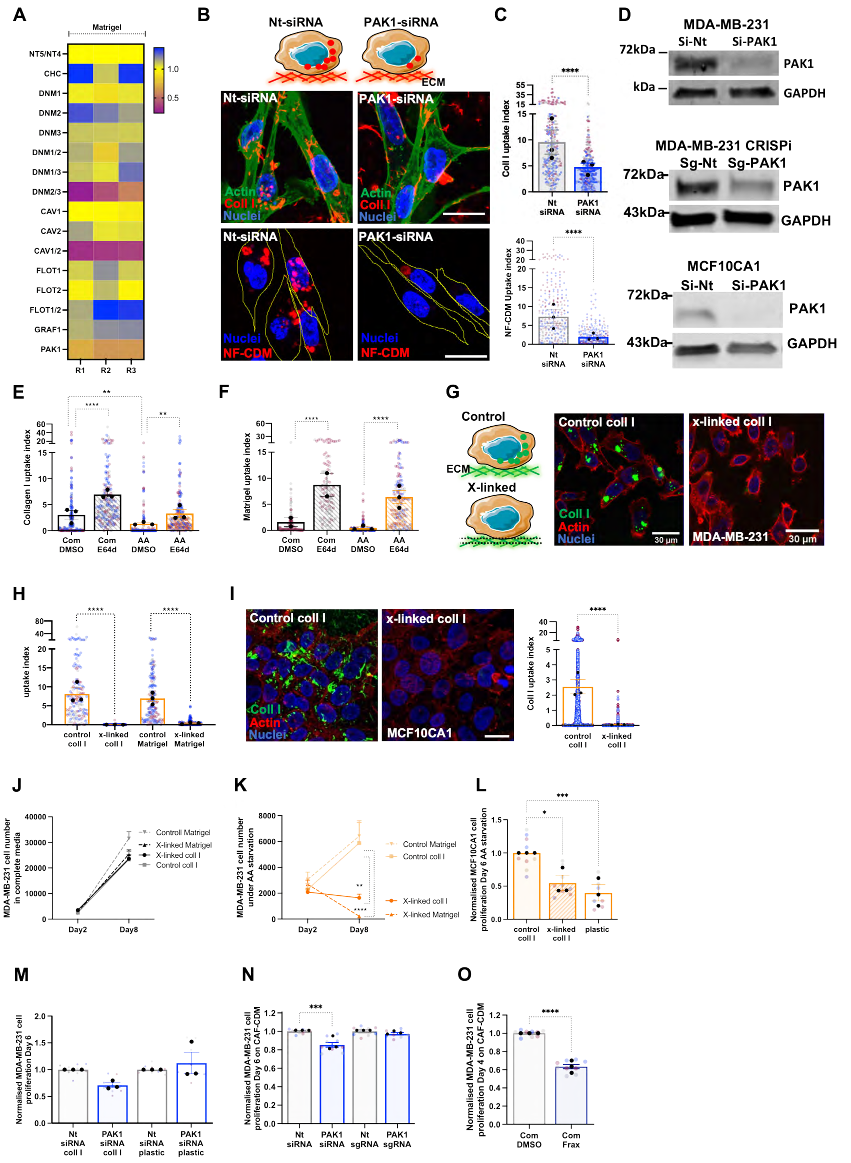


**Figure S4. ECM internalisation had limited effect on cell growth in complete media**. (**A**) MDA-MB-231 cells were transfected with siRNA targeting the indicated genes, seeded on pH-rodo labelled 0.5mg/ml Matrigel for 6 hrs, stained with Hoechst 33342, imaged live with an Opera Phenix miscroscope and analysed with Columbus software. (**B,C**) MDA-MB-231 cells were transfected with an siRNA targeting PAK1 (PAK1-siRNA) or a non-targeting siRNA control (nt-siRNA), plated on pH-rodo labelled NF-CDM (red) or Alexa Fluor 555 labelled 1mg/ml collagen I (coll I, red) for 6 hr, stained with Hoechst 33342 (blue) and imaged live with a Nikon A1 confocal microscope or fixed and stained for actin (green) and nuclei (blue). Scale bar, 20$\mu$m. ECM uptake index was quantified with Image J. (**D**) MDA-MB-231, MDA-MB-231 CRISPRi and MCF10CA1 cells were transfected with an siRNA targeting PAK1 (PAK1-siRNA), a non-targeting siRNA control (nt-siRNA), a synthetic guide RNA targeting PAK1 (PAK1-sgRNA) or a non-targeting synthetic guide RNA control (nt-sgRNA), lysed and PAK1 and GAPDH expression were measured by Western Blotting. MDA-MB-231 cells were plated under complete (Com) or amino acid depleted (AA) media on NHS-fluorescein labelled (**E**) 2mg/ml collagen I (coll I) or (**F**) 3mg/ml Matrigel coated dishes for 3 days, in the presence of the lysosomal inhibitor E64d (20$\mu$M) or DMSO (control). (**G-I**) 2mg/ml collagen I (coll I) or 3mg/ml Matrigel coated dishes were labelled with NHS-Fluorescein (green) and treated with 10% glutaraldehyde for 30 mins or left untreated. MDA-MB-231 (**G-H**) or MCF10CA1 (**I**) cells were plated on the cross-liked (x-linked) matrices for 3 days under amino acid (AA) starvation in the presence of E64d (20$\mu$M). Cells were fixed and stained for actin (red) and nuclei (blue). Samples were imaged with a Nikon A1 confocal microscope and ECM uptake index was calculated with image J. Bar, 30$\mu$m (**G**) and 20$\mu$m (**I**). More than 150 cells per condition in 3 independent experiments were analysed, the black dots represent the mean of individual experiments. **p<0.01, **** p<0.0001 Kruskal-Wallis, Dunn's multiple comparisons test. MDA-MB-231 (**J,K**) abd MCF10CA1 (**L**) cells were seeded on untreated (control) or cross-linked (X-linked) 2mg/ml collagen I (coll I) or 3mg/ml Matrigel under complete media (com, (**J**) or amino acid (AA) starvation (**K,L**) for 6 or 8 days. Cells were fixed and stained with Hoechst 33342. MDA-MB-231 (**M,N**) and MDA-MB-231 CRSPRi (**N**) cells were plated on 2mg/ml collagen I (coll I, **M**) or CAF-CDM (**N**), transfected with an siRNA targeting PAK1 (PAK1-siRNA), a non-targeting siRNA control (nt-siRNA, **M,N**), a synthetic guide RNA targeting PAK1 (PAK1-sgRNA) and a non-targeting synthetic guide RNA control (nt-sgRNA, **N**) and cultured in complete media for 6 days. Cells were fixed and stained with Hoechst 33342. Images were collected by ImageXpress micro and analysed by MetaXpress software. MDA-MB-231 cells (**O**) were grown on CAF-CDM in complete media for 4 days in the presence of 3$\mu$M FRAX597, fixed and stained with Hoechst 33342. Images were collected by ImageXpress micro and analysed by MetaXpress software. *p<0.05, *** p<0.001, **** p<0.0001 2way ANOVA, Tukey's multiple comparisons (**K**); Kruskal-Wallis, Dunn's multiple comparisons test (**N**); Mann-Whitney test (**O**).

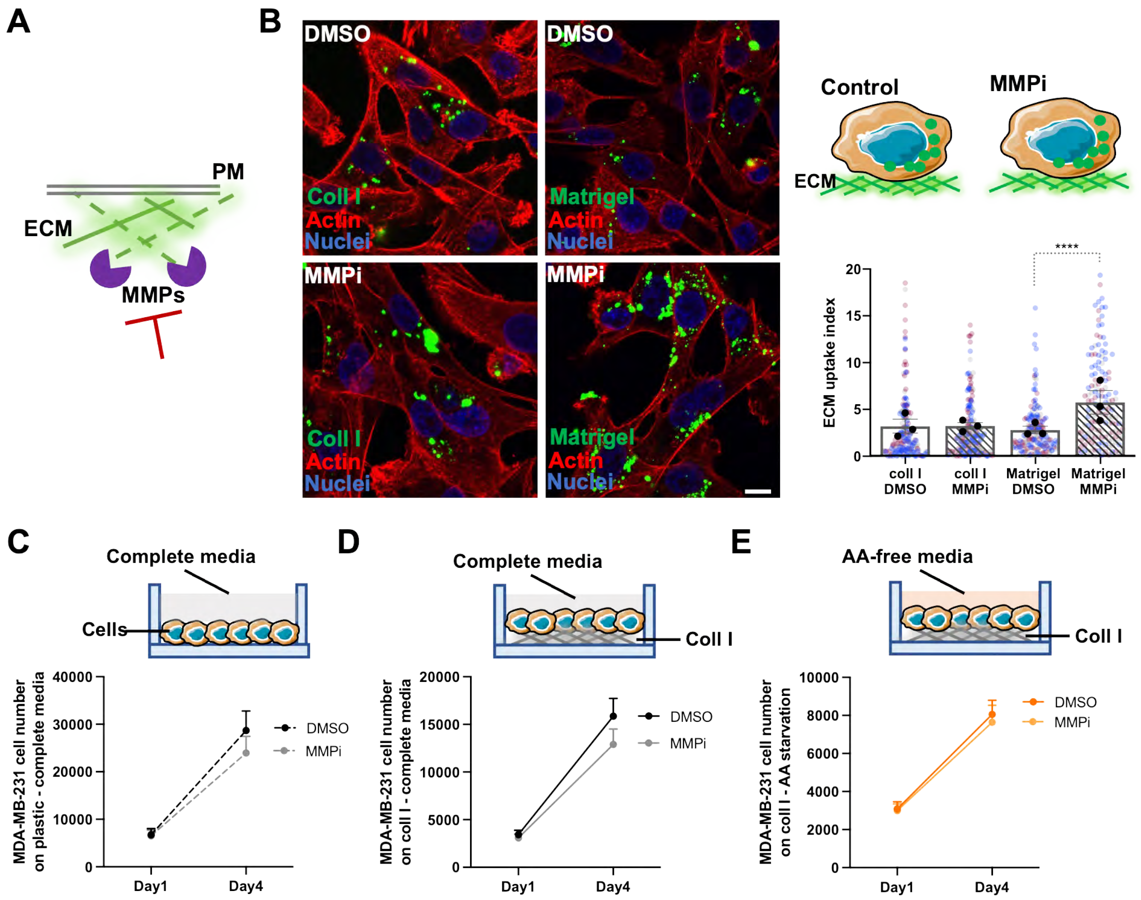


**Figure S5. MMP inhibition did not affect ECM uptake and cell growth**. (**A**) Schematic, MMP activity. (**B**) MDA-MB-231 cells were plated in complete media on NHS-fluorescein labelled 2mg/ml collagen I (coll I) or 3mg/ml Matrigel coated dishes for 3 days, in the presence of 20$\mu$M E64d and 10$\mu$M GM6001 (MMPi) or DMSO (control). Cells were fixed and stained for actin (red) and nuclei (blue). Samples were imaged with a Nikon A1 confocal microscope and ECM uptake index was calculated with image J. Bar, 10$\mu$m. About 150 cells per condition from 3 independent experiments were analysed, the black dots represent the mean of individual experiments. **** $p<0.0001$ Kruskal-Wallis, Dunn's multiple comparisons test. (**C-E**) MDA-MB-231 were seeded on (**C**) plastic or (**D,E**) 2mg/ml collagen I under complete (Com) or AA depleted (AA) media for 4 days. 10$\mu$M GM6001 (MMPi) or DMSO (control) were added to cells every 2 days. Cells were fixed and stained with Hoechst 33342. Images were collected by ImageXpress micro and analysed by MetaXpress software. Values are mean ± SEM from 3 independent experiments.

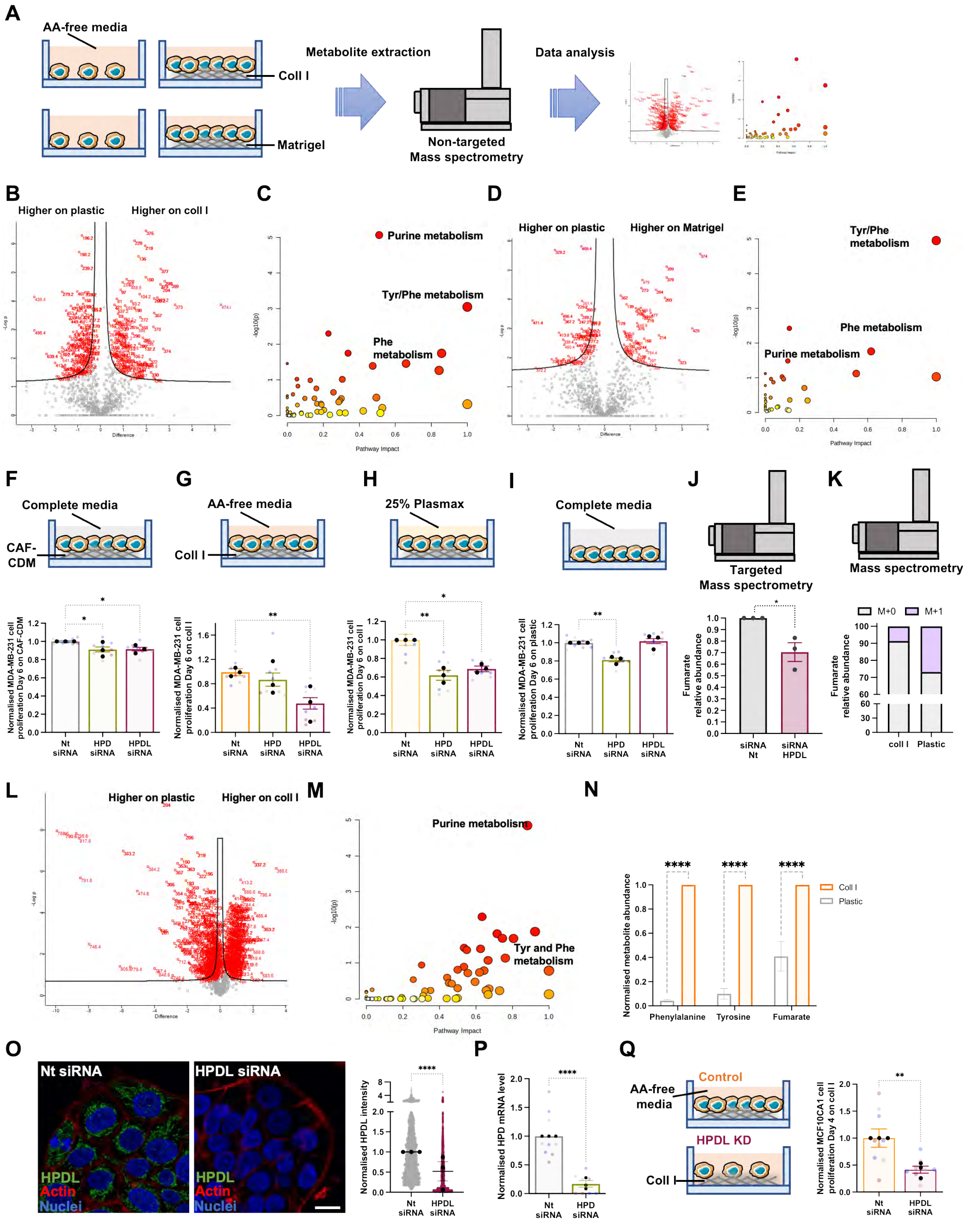

**Figure S6. Tyrosine catabolism did not affect cell growth on plastic.** MDA-MB-231 (**A-E**) and MCF10CA1 (**L,M**) cells were plated on plastic, 2mg/ml collagen I or 3mg/ml Matrigel for 6 days in amino acid-free media. Metabolites were extracted and quantified by non-targeted mass spectrometry. Volcano plots (**B,D,L**) and enriched metabolic pathways (**C,E,M**) are presented. (**F-I**) MDA-MB-231 or MCF10CA1 (**Q**) cells were plated on CAF-CDM (**F**), 2mg/ml collagen I (coll I, **G,P**) or plastic (**I**), transfected with siRNA targeting HPD (HPD siRNA), HPDL (HPDL siRNA) or non-targeting siRNA control (Nt siRNA) and cultured in complete, 25% Plasmax or amino acid-free media for 6 days. Cells were fixed and stained with Hoechst 33342. Images were collected by ImageXpress micro and analysed by MetaXpress software. Values are mean ± SEM and from three independent experiments. * $p<0.05$, ** $p<0.01$, **** $p<0.0001$ Kruskal-Wallis, Dunn's multiple comparisons test. (**J**) MDA-MB-231 cells were plated on 2mg/ml collagen I, transfected with siRNA targeting HPDL (HPDL siRNA) or non-targeting siRNA control (Nt siRNA) and cultured in amino acid depleted media for 3 days. Metabolites were extracted and fumarate was measured by targeted mass spectrometry. *$p=0.0126$ Mann-Whitney test. (**K**) MDA-MB-231 cells were grown in amino acid free media in the presence of C13 Tyrosine for 6 days. Metabolites were extracted and fumarate isotopologue abundance was quantified by targeted mass spectrometry. (**N**) Metabolites were prepared as in (A) and the levels of phenylalanine, tyrosine and fumarate were measured by targeted mass spectrometry. (**O**) MCF10CA1 cells transfected with siRNA targeting HPDL (HPDL siRNA) or non-targeting siRNA control (Nt siRNA), fixed and stained for HPDL (green), actin (red) and nuclei (blue). Images were collected with a Nikon A1 confocal microscope. Bar, 20$\mu$. HPDL intensity was quantified with ImageJ. (**P**) MDA-MB-231 cells transfected with siRNA targeting HPD (HPD siRNA) or non-targeting siRNA control (Nt siRNA), the mRNA was extracted and HPD expression was measured by qPCR. Images were collected with a Nikon A1 confocal microscope. Bar, 20$\mu$. HPDL intensity was quantified with ImageJ. Data are mean ± SEM from 3 independent experiments; **** $p<0.0001$, Mann-Whitney test.

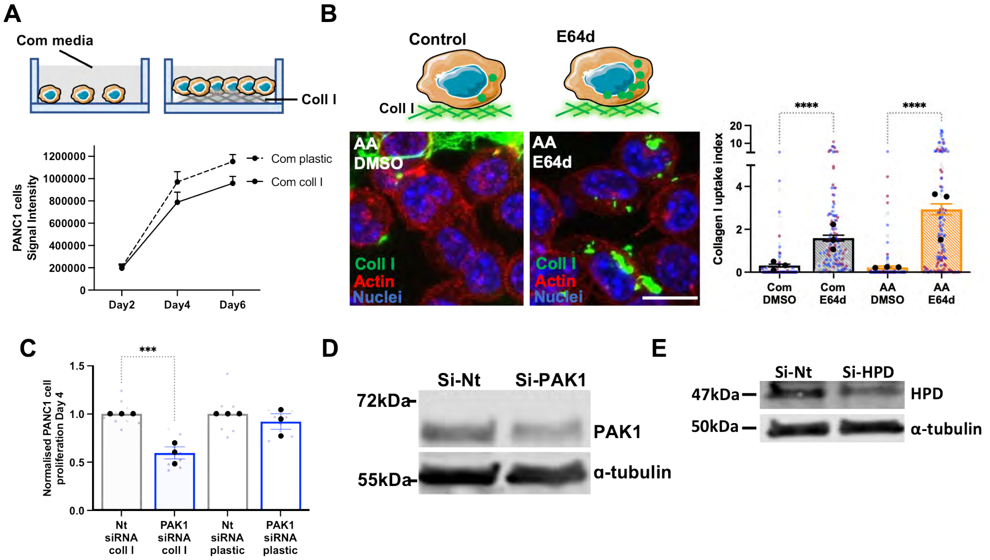


**Figure S7. PAK1 knock-down did not affect PDAC cell growth on plastic**. (**A**) PANC1 cells were seeded on plastic or 2mg/ml collagen I (coll I) for 6 days under complete media (com), fixed, stained with DRAQ5 and imaged with a Licor Odyssey system . Signal intensity was calculated by Image Studio Lite software. (**B**) PANC1 cells were plated under complete (Com) or amino acid depleted (AA) media on NHS-fluorescein labelled 2mg/ml collagen I (coll I) coated dishes for 3 days, in the presence of the lysosomal inhibitor E64d (20$\mu$M) or DMSO (control). Cells were fixed and stained for actin (red) and nuclei (blue). Samples were imaged with a Nikon A1 confocal microscope and ECM uptake index was calculated with image J. Bar, 20$\mu$m. About 200 cells per condition in three independent experiments were analysed, the black dots represent the mean of individual experiments. **** p<0.0001 Kruskal-Wallis, Dunn's multiple comparisons test. (**C**) PANC1 cells were plated on 2mg/ml collagen I (coll I) or plastic, transfected with an siRNA targeting PAK1 (PAK1-siRNA) or a non-targeting siRNA control (nt-siRNA) and cultured in complete media for 4 days. Cells were fixed and stained with Hoechst 33342. Images were collected by ImageXpress micro and analysed by MetaXpress software. Values are mean ± SEM and from 3 independent experiments (the black dots represent the mean of individual experiments). *** p<0.001 or Kruskal-Wallis, Dunn's multiple comparisons test. (**D,E**) PANC1 cells were plated transfected with an siRNA targeting PAK1 (PAK1-siRNA), an siRNA targeting HPD (HPD-siRNA) or a non-targeting siRNA control (nt-siRNA), lysed and analysed by Western Blotting.
